# Lipid metabolism links nutrient-exercise timing to insulin sensitivity in men classified as overweight or obese

**DOI:** 10.1101/742627

**Authors:** R.M. Edinburgh, H.E Bradley, N-F. Abdullah, S.L. Robinson, O.J. Chrzanowski-Smith, J-P. Walhin, S. Joanisse, K.N. Manolopoulos, A. Philp, A. Hengist, A. Chabowski, F.M. Brodsky, F. Koumanov, J.A. Betts, D. Thompson, G. A. Wallis, J.T. Gonzalez

## Abstract

**Context:** Pre-exercise nutrient availability alters acute metabolic responses to exercise, which could modulate training responsiveness. We hypothesised that in men with overweight/obesity, acute exercise before *versus* after nutrient ingestion would increase whole-body and intramuscular lipid utilization, translating into greater increases in oral glucose insulin sensitivity over 6-weeks of training.

**Design and Participants:** We showed in men with overweight/obesity (mean±SD for BMI: 30.2±3.5 kg×m^-2^ for acute, crossover study, 30.9±4.5 kg×m^-2^ for randomized, controlled, training study) a single exercise bout before *versus* after nutrient provision increased lipid utilisation at the whole-body level, but also in both type I (*p<*0.01) and type II muscle fibres (*p=*0.02). We then used a 6-week training intervention to show sustained, 2-fold increases in lipid utilisation with exercise before *versus* after nutrient provision (*p<*0.01).

**Main Outcome Measures:** Postprandial glycemia was not differentially affected by exercise training before *vs* after nutrient provision (*p>*0.05), yet plasma was reduced with exercise training before, but not after nutrient provision (*p=*0.03), resulting in increased oral glucose insulin sensitivity when training was performed before *versus* after nutrient provision (25±38 *vs* −21±32 mL×min^-1^×m^-2^; *p=*0.01) and this was associated with increased lipid utilisation during exercise (*r*=0.50, *p=*0.02). Regular exercise prior to nutrient provision augmented remodelling of skeletal muscle phospholipids and protein content of the glucose transport protein GLUT4 (*p<*0.05).

**Conclusions:** Experiments investigating exercise training and metabolic health should consider nutrient-exercise timing, and exercise performed before *versus* after nutrient intake (i.e., in the fasted state) may exert beneficial effects on lipid utilisation and reduce postprandial insulinemia.

**Précis:** Exercise in the fasted-*versus* fed-state increased intramuscular and whole-body lipid use, translating into increased muscle adaptation and insulin sensitivity when regularly performed over 6 weeks.

## INTRODUCTION

Postprandial hyperinsulinemia and associated peripheral insulin resistance are key drivers of metabolic diseases, such as type 2 diabetes (T2D) and cardiovascular disease (1–3). Obesity and a sedentary lifestyle are independently associated with changes in skeletal muscle that can reduce insulin sensitivity (4, 5) and increase hyperinsulinemia, contributing to elevated cardiovascular disease risk (2). Therefore, increasing insulin sensitivity and reducing postprandial insulinemia are important targets for interventions to reduce the risk of metabolic disease.

Regular exercise training represents a potent strategy to increase peripheral insulin sensitivity and to reduce postprandial insulinemia (6). The beneficial effects of exercise on oral glucose tolerance and insulin sensitivity can be attributed to both an ‘acute phase’ (during and straight after each bout of exercise performed) and the more enduring molecular adaptations that accrue in response to regular exercise (7). A single bout of endurance-type exercise activates contractile pathways in exercising muscle, which (independently of insulin) translocate the glucose transporter, GLUT4, to the plasma membrane and T-tubules to facilitate increased transmembrane glucose transport (8–10). The mechanisms that underlie the exercise-training induced increases in oral glucose insulin sensitivity include an increase in the total amount of time spent in the ‘acute phase’ (7), but also other adaptations that occur, such as changes in body composition (e.g. increased fat-free mass and reduced adiposity), an increased mitochondrial oxidative capacity (11), adaptations relating to glucose transport and insulin signaling pathways (12), and alterations to the lipid composition of skeletal muscle (13, 14).

Despite the potential for exercise to increase whole-body and peripheral insulin sensitivity, there can be substantial variability in the insulin sensitizing effects of fully-supervised exercise training programs (15). Crucially, this inter-individual variability for postprandial insulinemia following exercise training has also been shown to be greater than that of a control group (15), which demonstrates that some of this variability to exercise is true inter-individual variability (16). Nutritional status and thus the availability of metabolic substrates alters metabolism during and following exercise (17–20). Carbohydrate feeding before and during exercise suppresses whole-body and skeletal muscle lipid utilization (21, 22) and blunts the skeletal muscle mRNA expression of genes involved in exercise-adaptation for many hours post-exercise (23–25). This raises the possibility that nutrient-exercise interactions may regulate adaptive responses to exercise and thereby contribute to the apparent individual variability in exercise responsiveness *via* skeletal muscle adaptation and/or pathways relating to substrate metabolism.

Emerging data in lean, healthy men suggests that nutrient provision affects adaptive responses to exercise training (26, 27). However, feeding and fasting may exert different physiological responses in people who are overweight or obese compared to lean individuals, for example, extended morning fasting *versus* daily breakfast consumption upregulates the expression of genes involved in lipid turnover in adipose tissue in lean, but not in obese humans (28). Therefore, in order to fully understand the potential for nutrient-exercise timings to alter metabolism, exercise-adaptations and metabolic health in individuals at increased risk of metabolic disease, there is a need to study the most relevant populations, such as individuals classified as overweight or obese (29). It is currently unknown whether nutrient provision before *versus* after exercise affects adaptations to exercise training in these populations.

To this end, the aim of the present work was to assess the acute and chronic effects of manipulating nutrient-exercise timing on lipid metabolism, skeletal muscle adaptations, and oral glucose insulin sensitivity in men with overweight or obesity. We hypothesized that nutrient-exercise interactions would affect the acute metabolic responses to exercise, with increased whole-body and intramuscular lipid utilization with exercise before *versus* after nutrient provision. We also hypothesized that these acute responses to exercise before *versus* after nutrient provision would result in greater training-induced increases in oral glucose insulin sensitivity in men classified as overweight or obese.

## MATERIALS AND METHODS

### Ethical Approval

This project comprised two experiments. We first assessed the acute metabolic and mRNA responses to manipulating nutrient-exercise timing (**Acute Study**), followed by a 6-week randomized controlled trial to assess the longer-term (i.e. training) adaptations in response to nutrient-exercise timing (**Training Study**). All participants provided informed written consent prior to participation. Potential participants were excluded if they had any condition, or were taking any medication, known to alter any of the outcome measures. The studies were registered at https://clinicaltrials.gov (NCT02397304 and NCT02744183, respectively). Protocols were approved by the National Health Service Research Ethics Committee (15/WM/0128 & 16/SW/0260, respectively) and experiments were conducted in accordance with the Declaration of Helsinki.

### Acute Study

In the **Acute Study**, 12 sedentary, men classified as overweight or obese were recruited from the Birmingham region of the UK. The main exclusion criteria included being regularly physically active, having hypertension or possible (undiagnosed) T2D. Participant characteristics are shown in Table 1.

**Table 1.**
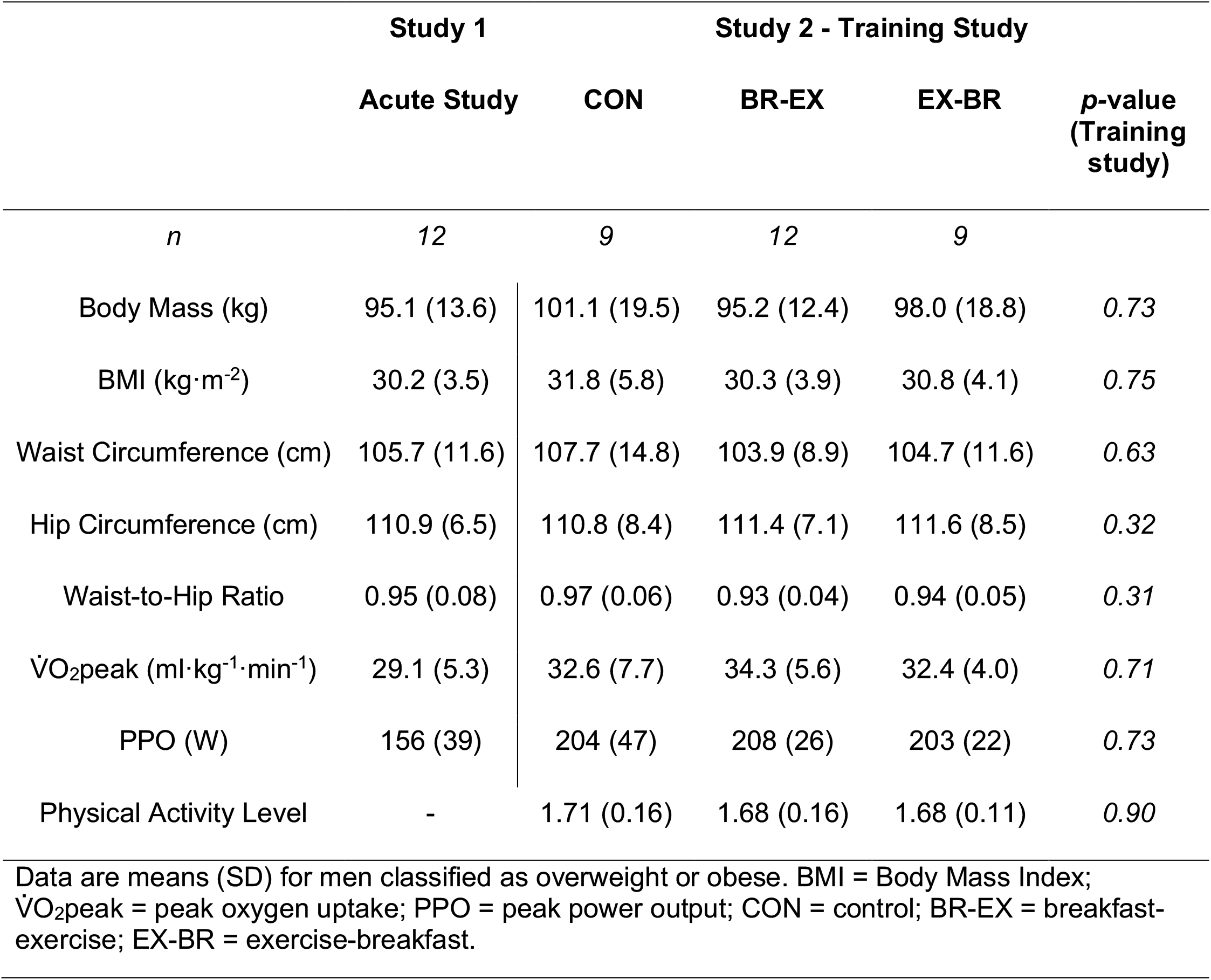
Participant characteristics

|  | Study 1 |  | Study 2 - Training Study |  | *p*-value<br>(Training<br>study) |
| --- | --- | --- | --- | --- | --- |
|  | Acute Study | CON | BR-EX | EX-BR |  |
| *n* | 12 | 9 | 12 | 9 |  |
| Body Mass (kg) | 95.1 (13.6) | 101.1 (19.5) | 95.2 (12.4) | 98.0 (18.8) | 0.73 |
| BMI (kg·m<sup>-2</sup>) | 30.2 (3.5) | 31.8 (5.8) | 30.3 (3.9) | 30.8 (4.1) | 0.75 |
| Waist Circumference (cm) | 105.7 (11.6) | 107.7 (14.8) | 103.9 (8.9) | 104.7 (11.6) | 0.63 |
| Hip Circumference (cm) | 110.9 (6.5) | 110.8 (8.4) | 111.4 (7.1) | 111.6 (8.5) | 0.32 |
| Waist-to-Hip Ratio | 0.95 (0.08) | 0.97 (0.06) | 0.93 (0.04) | 0.94 (0.05) | 0.31 |
| VO<sub>2</sub>peak (ml·kg<sup>-1</sup>·min<sup>-1</sup>) | 29.1 (5.3) | 32.6 (7.7) | 34.3 (5.6) | 32.4 (4.0) | 0.71 |
| PPO (W) | 156 (39) | 204 (47) | 208 (26) | 203 (22) | 0.73 |
| Physical Activity Level | - | 1.71 (0.16) | 1.68 (0.16) | 1.68 (0.11) | 0.90 |

Data are means (SD) for men classified as overweight or obese. BMI = Body Mass Index; VO<sub>2</sub>peak = peak oxygen uptake; PPO = peak power output; CON = control; BR-EX = breakfast-exercise; EX-BR = exercise-breakfast.

This was a randomised cross-over study where on one visit (breakfast-exercise or BR-EX), a standardised breakfast (cornflake cereal with skimmed milk, wholemeal toast, sunflower spread and strawberry jam) was consumed upon arrival at the laboratory (and following 48 h of diet control). The breakfast provided 25% of estimated daily energy requirements [calculated as resting metabolic rate (RMR) multiplied by a physical activity factor of 1.53 (30)] and was 65% carbohydrate, 20% fat and 15% protein. After a 90-min period of rest, 60 min of cycling exercise was then performed at 65% peak oxygen uptake (V̇O_2_ peak). Expired gas samples were collected at 25-30 min and 55-60 min of exercise to determine whole-body substrate utilisation rates. Blood was sampled in the overnight-fasted state, at 45 min post breakfast and immediately before exercise was performed (90 min post breakfast), every 30 min during exercise and at 60 min intervals during a 3-h post-exercise recovery. In a subset of participants (*n*=8) *vastus lateralis* muscle was sampled pre-and immediately post-exercise to assess fiber-type specific intramuscular triglyceride (IMTG) and mixed-muscle glycogen utilization. A third muscle sample (taken at 3 h post-exercise) was used to assess the intramuscular gene expression (mRNA) responses to exercise (*n*=7). On the other visit (exercise-breakfast or EX-BR) the participants completed the same protocol, but the breakfast was consumed immediately after the post-exercise muscle sample. The primary outcome for the **Acute Study** was intramuscular lipid utilisation during exercise performed before *versus* after nutrient ingestion.

### Training Study

To assess longer-term adaptive (i.e. training) responses to altering nutrient-exercise timing (**Training Study**) we recruited 30, overweight and obese, sedentary men (self-reported non-exercisers) from the Bath region of the UK (Table 1). This was a single-blind, randomized, controlled trial, with participants allocated to a no-exercise control group (CON; *n*=9) a breakfast before exercise group (BR-EX; *n*=12) or an exercise before breakfast group (EX-BR; *n*=9) for 6-weeks (Figure 1). The exercise was supervised moderate-intensity cycling (Monark Exercise AB, Vansbro, Sweden) performed 3 times per week, starting at 50% peak power output [PPO] (weeks 1-3) and increasing to 55% PPO (weeks 4-6). The duration of the exercise sessions progressed from 30-(week 1) to 40-(week 2) to 50-min (weeks 3-6). All sessions were supervised at the University of Bath. During every one of the 336 exercise training sessions, 1-min expired gas samples were collected every 10 min to assess substrate utilization and heart rate (Polar Electro Oy, Kempele, Finland) and ratings of perceived exertion (31) were recorded.

**Figure 1.**
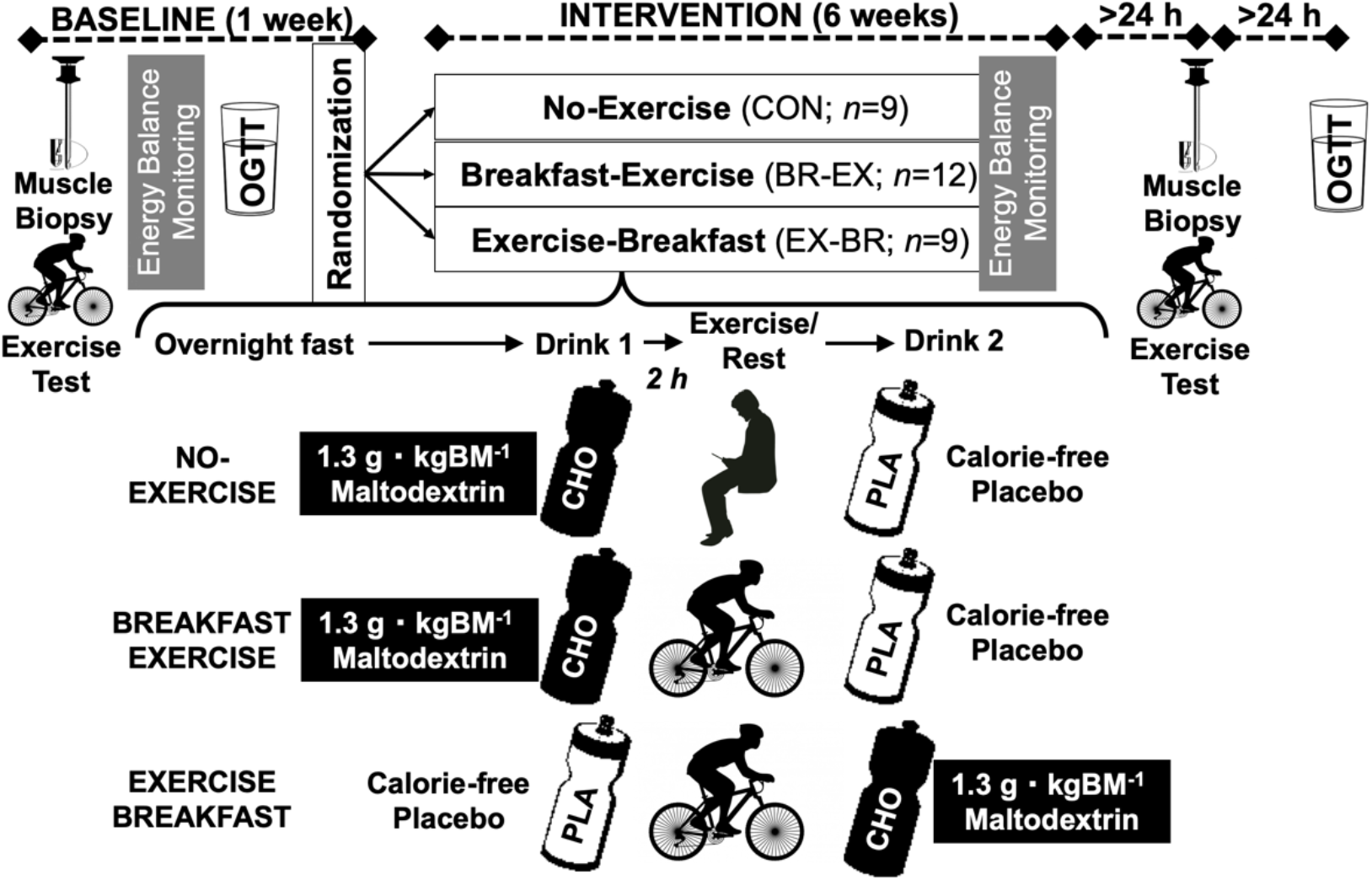
Protocol schematic for the training study.

Participants ate their evening meal before 2000 h the evening prior to any exercise sessions. Participants in BR-EX were given a drink in an opaque bottle made from 1.3 g carbohydrate·kg body mass^-1^ maltodextrin (MyProtein, Northwich, UK) with vanilla flavoring (20% carbohydrate solution) for consumption 2-h before exercise. They were asked not to eat or drink anything else (except water *ad libitum*) in this period and confirmed they had consumed the drink before exercising. After exercise, they were provided a taste matched placebo (water and vanilla flavoring) to consume 2-h after exercise and were asked not to consume anything else during this period. Participants in EX-BR were given the same drinks, but with the order of the drinks reversed. Participants in CON were given the same drinks for three days per week during the intervention, with the carbohydrate drink as breakfast (0800-0900 h) and the placebo for consumption with their lunch (1100-1300 h). These participants were asked not to consume anything else between the drinks. There were no other diet controls in the intervention. Blinding of the groups was deemed successful, because at exit interview, 25 participants (83%) revealed they could not detect a difference between the carbohydrate and placebo drinks or could not identify which contained carbohydrate. Five participants determined which drink had carbohydrate (CON *n*=1, BR-EX *n*=2, EX-BR *n*=1), but this is within the proportion that could do so at random.

Pre-and post-intervention, an oral glucose tolerance test (OGTT), a *vastus lateralis* muscle sample (fasting, rested state) and an exercise test (to assess V̇O_2_ peak and the capacity for lipid utilization during exercise in the fasted-state) were undertaken. Post-intervention tests were between 24 h to 48 h (for muscle sampling) and 48 h to 72 h (for OGTT) after the last exercise training session, to reduce any residual effects of the last exercise bout performed on these measurements. The primary outcome for the **Training Study** was the pre-to post-intervention change in the glycaemic and insulinemic responses to the OGTT, which were also used to derive an index of oral glucose insulin sensitivity (as described subsequently).

### Pre-trial standardizations

For both studies, the participants were asked to maintain their normal physical activity behaviors and to abstain from alcoholic and caffeinated drinks for 24 h prior to all main laboratory trials. Food intake ceased at 2000 h ± 1 h on the evening before testing and participants fasted overnight (minimum of 10 h). For all trials, participants arrived at the laboratory at 0800 ± 1 h, with the exact time replicated for subsequent trials. For the **Acute Study**, participants were provided with a standardized weight maintaining diet (50% carbohydrate, 35% fat, 15% protein) based on their estimated energy requirements (RMR multiplied by the physical activity factor of 1.53 as stated previously) for consumption for 48 h prior to main trials. For the **Training Study**, they recorded the composition of their evening meal on the day before a pre-intervention trial and replicated this meal for the post-intervention trial, in line with guidelines for testing postprandial glycemic control (32). We have shown that this protocol produces fasting muscle and liver glycogen and fasting intramuscular lipid concentrations that are consistent across trial days (33).

### Anthropometry

Stature was measured to the nearest 0.1 cm using a stadiometer (Seca Ltd, Birmingham, UK). Body mass was measured to the nearest 0.1 kg using electronic weighing scales (**Acute Study**: OHaus Champ II Scales, USA; **Training Study**: BC543 Monitor, Tanita, Japan). Waist and hip circumferences were measured to the nearest 0.1 cm and according to the World Health Organization guidelines.

### Exercise tests

Participants completed exercise tests on an electronically-braked ergometer. In the **Acute Study**, the starting intensity was 35 W, and this was increased by 35 W every 3 min until volitional exhaustion. In the **Training Study**, the starting intensity for the exercise test was 50 W which was increased by 25 W every 3 min. Heart rate (Polar Electro Oy, Kempele, Finland) and continuous breath-by-breath measurements were recorded (**Acute Study**: Oxycon Pro, Jaeger, Wurzburg, Germany; **Training Study**: TrueOne2400, ParvoMedics, Sandy, USA). Volume and gas analyzers were calibrated using a 3-L calibration syringe (Hans Rudolph, Kansas City, USA) and a calibration gas (16.04% O_2_, 5.06% CO_2_; BOC Industrial Gases, Linde AG, Germany). Peak power output (PPO) was calculated as the work rate of the final completed stage, plus the fraction of time in the final non-completed stage, multiplied by the W increment. V̇O_2_ peak was the highest measured V̇O_2_ over a 30 s period, using methods and attainment criteria previously reported (34).

### Blood sampling and analysis

In the **Acute Study**, 10 mL blood was sampled from an antecubital forearm vein and 6 mL was dispensed into ethylenediaminetetraacetic acid-coated tubes (BD, Oxford, UK) and centrifuged (4°C at 3500 rpm) for 15 min (Heraeus Biofuge Primo R, Kendro Laboratory Products Plc., UK). Resultant plasma was dispensed into 0.5 mL aliquots and frozen at −20°C, before longer-term storage at −80°C. A proportion of the sample (4 mL) was allowed to clot in a plain vacutainer prior to centrifugation, for serum. Samples were analyzed for plasma glucose, glycerol and NEFA using an ILAB 650 Clinical Chemistry Analyzer (Instrumentation Laboratory, Warrington, UK). Serum insulin concentrations were measured with an ELISA kit (Invitrogen; Cat#KAQ1251) and Biotek ELx800 analyzer (Biotek Instruments, Vermont, USA).

In the **Training Study**, prior to blood sampling participants placed their dominant hand into a heated-air box set to 55°C. After 15 min of rest, a catheter was placed (retrograde) into a dorsal hand vein and 10-mL of arterialized blood was drawn for a baseline sample in the overnight-fasted state (35). Then a 75-g OGTT was completed and arterialized blood sampled every 15 min for 2 h and processed (as detailed above) for plasma. Plasma glucose (intra-assay CV: 2.50%), glycerol, triglyceride (glycerol-blanked), and total-HDL-and LDL-cholesterol concentrations were measured using an automated analyzer (Daytona; Randox Lab, Crumlin, UK). Plasma insulin (Mercodia AB; reference #10-1113-01) and C-peptide (Sigma Aldrich; reference #EZHCP-20K) concentrations were measured using commercially available ELISA kits (intra-assay CV for insulin: 3.86% and for C-peptide: 4.26%). Non-esterified fatty acid (NEFA) concentrations were assessed via an enzymatic colorimetric kit (WAKO Diagnostics; references #999-34691/#991-34891; intra-assay CV: 7.95%). All analysis was done in batch and for a given participant all samples were included on the same plate.

### Muscle sampling

All *vastus lateralis* skeletal muscle samples were collected under local anesthesia (∼ 5 mL 1% lidocaine, Hameln Pharmaceuticals Ltd., Brockworth, UK) and from a 3-6-mm incision at the anterior aspect of the thigh using a 5-mm Bergstrom biopsy needle technique adapted for suction. For the **Acute Study**, samples were collected pre-and immediately post-exercise and at 3 h post-exercise. To enable the analysis of the IMTG content ∼ 15-20 mg of each sample was embedded in Tissue-Tek OCT (Sigma Aldrich, Dorset, UK) on cork disc and frozen in liquid nitrogen cooled isopentane, before being transferred into an aluminium cryotube and stored at −80°C. Remaining muscle (for glycogen and gene expression analysis) was frozen in liquid nitrogen and stored at −80°C. For the **Training Study**, samples were collected pre-and post-intervention with participants in a fasted, resting state, with both samples collected from their dominant leg. Muscle was extracted from the needle and frozen in liquid nitrogen, before storage at −80 °C. Frozen wet muscle (80-100 mg) was freeze-dried and powdered, with visible blood and connective tissue removed. Ice cold lysis buffer (50 mM Tris [pH 7.4], 150 mM NaCl, 0.5% Sodium deoxycholate; 0.1% SDS and 0.1% NP-40) with protease and phosphatase inhibitors was added. Samples were homogenized with a dounce homogenizer, before a 60 min incubation (4°C with rotation) and 10 min centrifugation (4°C and 20,000 g). The protein content of the resultant supernatant was measured using a bicinchoninic acid assay.

### Intramuscular triglyceride (Acute Study)

The muscle mounted in Tissue-Tek was cut into 5 μm thick transverse sections with a cryostat at −25°C (Bright 5040, Bright Instrument Company; Huntingdon, England) and collected onto an uncoated glass slide and frozen immediately after sectioning. Each slide had 4 samples for a participant (pre- and post-exercise both trials) to decrease variation in staining intensity between muscle sections and slides were prepared and analyzed for each participant. For analysis, cryosections were removed from the freezer and fixed immediately in 3.7% formaldehyde for 60 min. Slides were then rinsed with distilled water (3 x 30 s) and treated for 5 min with 0.5% Triton-X100 in phosphate-buffered solution (PBS; 137 mmol·L^-1^ sodium chloride, 3 mmol·L^-1^ potassium chloride, 8 mmol·L^-1^ sodium phosphate dibasic, 3 mmol·L^-1^ potassium phosphate monobasic). The slides were washed (3 x 5 min in PBS) and incubated for 2 h at room temperature with anti-myosin heavy chain I antibody (MHCI; mouse IgM, Developmental Studies Hybridoma Bank: reference #A4.480) and anti-dystrophin antibody (mouse IgG2b, Sigma Aldrich: reference *#*D8168) in 5% goat serum diluted in PBS (1:1 PBS dilution). This was followed by washes in PBS (3 x 5 min), after which conjugated secondary antibodies [goat anti mouse (GAM) IgM conjugated to AlexaFluor 633 for MHCI; Thermo Fisher: reference *#*A21046; and GAM IgG2b conjugated to AlexaFluor 594 for dystrophin; Thermo Fisher: reference *#*A21145] were added and incubated at room temperature (30 min) followed by washes in PBS. Then, muscle sections were incubated in BODIPY 493/503 solution (Thermo Fisher: reference #D3922) for 20 min at room temperature in a dark room before washes (2 x 3 min in PBS). Stained sections were embedded in Mowiol 4-88 mounting medium (Fluka: reference #81381) and covered with a coverslip. Slides were left to dry overnight at room temperature before analysis by confocal microscope in duplicate (DMIRE2, Leica Microsystems; 40x oil objective; 1.25 NA). An argon laser 488 nm was used to excite BODIPY-493/503 (emission 510-652 nm), while a helium-neon 594 nm and 633 nm laser line were used to excite Alexa Fluor 594 (dystrophin, emission 6680698 nm) and AlexaFluor 633 (MHCI, emission 698-808 nm), respectively. Images were scanned in projection of 4 lines in 1024×1025 pixels format. Quantification of the lipid droplets was performed using Image J software and the intramuscular triglyceride (IMTG) content of each sample was calculated as the percent area of bodipy staining of the total fiber area ([BODIPY stained area [um^2^] / area of muscle [um^2^]*100).

### Muscle glycogen (Acute Study)

Muscle glycogen concentrations were measured using a method described previously (36). Briefly, 10-15 mg of frozen tissue was powdered and transferred into a glass tube pre-cooled on dry ice. Thereafter, the samples were hydrolyzed by adding a 500 µl of 2M HCL and then incubated for 2 h at 95 °C. After cooling to room temperature, 500µl 2M NaOH was added. Samples were centrifuged and the supernatant was analyzed for glucose concentrations using an ILAB 650 Clinical Chemistry Analyzer (Instrumentation Laboratory, Warrington, UK).

### Gene expression (Acute Study)

The mRNA expression of 34 metabolic genes was analyzed using a custom RT2 Profiler PCR Array (Qiagen, USA). First, RNA was extracted from 20-40 mg of powdered muscle tissue using Tri reagent (1 mL, Sigma Aldrich, UK, T9424). After addition of chloroform (200 uL, Acros organics 268320025), tubes were incubated at room temperature for 5 min and centrifuged for 10 min (4 °C at 12 000 g). The RNA phase was mixed with an equal volume of ice cold 70% ethanol and RNA was purified on Reliaprep spin columns (Promega, USA, Z6111) as per manufacturer’s instructions. The LVis function of the FLUOstar Omega microplate reader was used to measure RNA concentrations to ensure all samples for each participant had the same amount of RNA (184 ng - 400 ng) and samples were reverse transcribed to cDNA using the RT2 First Strand kit (Qiagen, UK, 330401). Quantitative RT-PCR analysis was performed using custom designed 384-well RT2 PCR Profiler Arrays (Qiagen) and RT2 SYBR Green Mastermix (Qiagen) on a CFX384 Real-Time PCR Detection system (BioRad). 2.8 ng cDNA was added to each well. All primers that were used are commercially available (Table 2). The absence of genomic DNA, the efficiency of reverse-transcription and the efficiency of the PCR assay were assessed for each sample and conformed to manufacturer’s limits. Relative mRNA expression was determined via the 2^-ΔΔCT^ method (37). Housekeeper genes (*β*actin [Refseq# NM_001101]; ribosomal protein lateral stalk subunit P0 [Refseq# NM_001002] and *β*-2-microglobulin [Refseq# NM_004048] were internal controls.

**Table 2.** List of genes analyzed for mRNA expression

| **Gene** | **Qiagen catalogue number** | **Refseq#** |
| --- | --- | --- |
| *CD36* | Cat#PPH01356A | NM_000072 |
| *SLC27A1* | Cat#PPH17902A | NM_198580 |
| *SLC27A4* | Cat#PPH00471A | NM_005094 |
| *FABP3* | Cat#PPH02460C | NM_004102 |
| *FABP4* | Cat#PPH02382F | NM_001442 |
| *ACSL1* | Cat#PPH19272A | NM_001995 |
| *ACSL6* | Cat#PPH08013A | NM_001009185 |
| *CPT1B* | Cat#PPH20905B | NM_001145134 |
| *CPT2* | Cat#PPH15572A | NM_000098 |
| *ACACA* | Cat#PPH02316A | NM_000664 |
| *ACACB* | Cat#PPH02301A | NM_001093 |
| *MLYCD* | Cat#PPH12795A | NM_012213 |
| *HADHA* | Cat#PPH10000B | NM_000182 |
| *GPAM* | Cat#PPH06361A | NM_001244949 |
| *DGAT1* | Cat#PPH23420F | NM_012079 |
| *PNPLA2* | Cat#PPH11403B | NM_020376 |
| *LIPE* | Cat#PPH02383A | NM_005357 |
| *PDK4* | Cat#PPH07615A | NM_002612 |
| *PDK2* | Cat#PPH00810A | NM_001199898 |
| *GYG1* | Cat#PPH13614A | NM_001184720 |
| *GYS1* | Cat#PPH00988C | NM_001161587 |
| *PRKAA1* | Cat#PPH00043B | NM_206907 |
| *PRKAA2* | Cat#PPH15207A | NM_006252 |
| *PRKAB2* | Cat#PPH09415B | NM_005399 |
| *PRKAG1* | Cat#PPH07190A | NM_001206709 |
| *PPARGC1A* | Cat#PPH00461F | NM_013261 |
| *PPARA* | Cat#PPH01281B | NM_001001928 |
| *PPARD* | Cat#PPH00455A | NM_001171818 |
| *UCP3* | Cat#PPH06066A | NM_003356 |
| *IL6* | Cat#PPH00560C | NM_000600 |
| *SLC2A4* | Cat#PPH02326A | NM_001042 |
| *IRS1* | Cat#PPH02328A | NM_005544 |
| *IRS2* | Cat#PPH02297A | NM_003749 |
| *AKT2* | Cat#PPH00289F | NM_001243027 |

### Western blotting (Training Study)

For western blots, 40 µg of protein was loaded for each sample and separated via sodium dodecyl sulfate polyacrylamide gel electrophoresis on Tris-glycine SDS– polyacrylamide gels (15% for OXPHOS and CPT-1, 10% CD36 and GLUT4 and 8% for AMPK, CHC22, CHC17, AkT and AS160). Gels were electro-blotted (semi-dry transfer) onto a nitrocellulose membrane and were then washed in Tris-buffered saline (0.09% NaCl, 100 mM Tris–HCl pH 7.4) with 0.1% Tween 20 (TBS-T) and incubated for 30 min in a blocking solution (5% non-fat milk in TBS-T). Membranes were incubated overnight at 4 °C with primary antibodies against OXPHOS (Abcam: reference #ab110411), CPT-1 (Abcam: reference #ab134988), CD36 (Abcam: reference #ab133625), GLUT4 [self-raised rabbit polyclonal antibody against the C-terminus of GLUT4 (38)], CHC22 [SHL-KS, affinity purified self-raised rabbit polyclonal against the CHC22 C-terminus cross-absorbed against the CHC17 C-terminus (39)], CHC17 [TD.1 self-raised mouse monoclonal against CHC17 terminal domain (40)] AMPKα (Cell Signalling Technologies: reference #2532), Akt (Cell Signaling Technologies: reference #3063), AS160 (Millipore: reference #07-741). It should be noted that the AMPKα antibody recognises both α1 and AMPKα2 isoforms of the catalytic subunit and does not detect the regulatory AMPKβ or AMPKγ subunits (41). Following incubation with the primary antibodies, the membranes were washed in TBS-T and incubated for 60 min in a 1:4000 dilution of anti-species IgG horseradish peroxidase-conjugated secondary antibodies in the aforementioned blocking solution. After further washes, membranes were incubated in an enhanced chemiluminescence reagent and visualized (EpiChemi II Darkroom, UVP, Upland, USA). The band densities were quantified using Image Studio Lite software (Version 5.2; LI-COR, Nebraska, USA) and were normalized to either GAPDH (Proteintech: reference #60004-1-Ig) or Actin (Sigma Aldrich: reference #A2066), before the pre-to post-intervention change was calculated. Pre-and post-intervention samples from any given participant were included on the same gel. Citrate synthase activity was measured using a commercially available assay (Abcam: reference #ab119692).

### Phospholipid composition (Training Study)

Samples were freeze-dried, powdered under liquid nitrogen, and transferred into glass tubes containing 2 ml of methanol and butylated hydroxytoluene (0.01%) and heptadecanoic acid (as an internal standard), followed by the addition of 4 mL of chloroform and 1.5 mL of water, before lipids were extracted. The lipid containing fraction was transferred into thin-layer chromatography (TLC; Kieselgel 60, 0.22 mm, Merck, Darmstadt, Germany) silica plates and lipids were separated by TLC with a heptane: isopropyl ether: acetic acid (60:40:3, vol/vol/vol) resolving solution. Lipid bands were made visible by spraying the plates with a 0.2% solution of 3’7’-dichlorofluorescin in methanol and recognized under ultraviolet light using standards on the plates. Then the gel bands containing phospholipids were scraped off the plates, transferred into screw cap tubes and transmethylated with BF3/methanol. The fatty acid methyl esters (FAMEs) were then dissolved in hexane and analyzed by GLC. A Hewlett-Packard 5890 Series II gas chromatograph with Varian CP-SIL capillary column (100 m, internal diameter of 0.25 mm) and flame-ionization detector were used. In accordance with the retention times of standards, the individual long-chain fatty acids quantification was performed. The content of phospholipids was estimated as the sum of the total fatty acid species and expressed in nanomoles per milligram of dry mass (42, 43).

### Energy expenditure and intake (Training Study)

Average daily energy expenditure was calculated as the sum of the resting metabolic rate (RMR), diet-induced thermogenesis (10% of self-reported daily energy intake) and physical activity energy expenditure (PAEE). To assess RMR, participants rested in a semi-supine position for 15 min before 4 x 5-min expired air samples were collected (44). The participants were provided with the mouthpiece 1 min prior to sample collections (as a stabilization period) which were collected into a 200-L Douglas bag (Hans Rudolph, Kansas City, USA) via falconia tubing (Baxter, Woodhouse and Taylor Ltd, Macclesfield, UK). Concurrent measures of inspired air were also made to correct for changes in the ambient O_2_ and CO_2_ concentrations. Expired O_2_ and CO_2_ concentrations were measured in a volume of each sample using paramagnetic and infrared transducers (Mini HF 5200, Servomex Group Ltd., Crowborough, UK). The sensor was calibrated with low (0% O_2_ and 0% CO_2_) and high (16.04% O_2_, 5.06% CO_2_) calibration gases (BOC Industrial Gases, Munich, Germany). Substrate utilization rates were then calculated via stochiometric equations (45, 46). Energy expenditure was calculated assuming that fatty acids, glucose and glycogen provide 40.81 kJ·g^-1^, 15.64 kJ·g^-1^ and 17.36 kJ·g^-1^ of energy, respectively. To measure free-living PAEE, participants wore an Actiheart^TM^ monitor over 7 days (Cambridge Neurotechnology, Papworth, UK). This monitor integrates accelerometry and heart rate signals and has been validated as a measure of energy expenditure (47–49). Energy expenditure and heart rate values from rest and exercise were entered in the Actiheart^TM^ software for an individually calibrated model. Participants were also asked to keep a written record of their food and fluid intake for 4 days over a typical 7-day period (including a weekend day) pre-and during the last week of the intervention. Weighing scales were provided to increase the accuracy of records. Records were analyzed using Nutritics software (Nutritics Ltd., Dublin, Ireland). The macronutrient composition of each food was taken from the manufacturer’s labels, but if this was not possible (e.g. fresh products) foods were analyzed via the software database or comparable brands were used to provide the relevant information, and this was kept constant across records.

### Statistics

In the **Acute Study**, the sample size was based upon data demonstrating an attenuation of intramuscular lipid utilization during exercise with carbohydrate intake before and during exercise with an effect size of *d* = 1.5. (22) We aimed to recruit 12 participants assuming at least 8 participants would complete the study with biopsies to provide >90% power with α set at 0.05. In the **Training Study**, a sample size estimation was completed using data from a training study in healthy, lean men (26). In that study, a change in the plasma glucose AUC for an oral glucose tolerance test (OGTT) of −65±53 mmol·min·L^-1^ was shown in an EX-BR group *versus* +21±47 mmol·min·L^-1^ for a CON group. With α set at 0.05, 9 participants were required for a >90% chance of detecting this effect. We therefore recruited 30 participants to account for the possibility of an unequal allocation of participants across three groups when using a stratified randomization schedule. Participants were allocated to the CON (*n*=9), BR-EX (*n*=12) or EX-BR (*n*=9) groups using this schedule, which was generated by JPW and included a factor for Physical Activity Level (PAL) and the time-averaged glucose AUC for the baseline OGTT which was assessed using a Freestyle Freedom Lite Glucose Meter. This was to ensure an even distribution of less (PAL <1.65) and more active (PAL >1.65) participants and participants with glucose AUC values above or below 8 mmol·L^-1^.

Data are presented as means ± 95% confidence intervals (CI), except for participant characteristics (which are mean ± SD). A Shapiro-Wilk test was performed to test for normal distribution and if this was not obtained, non-parametric tests (e.g. Wilcoxon matched-pairs signed rank tests) were employed. In the **Acute Study**, differences between groups were assessed with paired *t-*tests or a two-way repeated measures ANOVA (for variables dependent on time). In the **Training Study**, one-way ANOVAs were used to assess differences between groups at baseline and two-way mixed-design ANOVAs were used to assess differences between groups in response to the intervention (group x time). If interaction effects were identified, independent *t*-tests were used to locate variance, with Holm-Bonferroni step-wise adjustments made. Correlations between variables were explored using Pearson *r* or Spearman R for normal or non-normal distributions, respectively. A significance level of *p<*0.05 was always used. The area under the concentration-time curve (AUC) was calculated via the trapezoid rule and divided by the duration of an observation period of interest for a time-averaged summary value. Plasma glucose and insulin concentrations were used to assess oral glucose insulin sensitivity (the OGIS index) as per instructions provided at; http://webmet.pd.cnr.it/ogis/ (50). Statistical analyses were completed on IBM SPSS statistics V 22 for windows (except for the Holm-Bonferroni adjustments which were completed on Microsoft Excel) and Graph Pad Prism 7 was used to prepare the figures. As we were unable to collect data from all participants for all measured outcomes the *n* are always displayed in all figure and table captions.

## RESULTS

**Exercise before nutrient ingestion increases whole-body and skeletal muscle lipid utilization but does not differentially modulate muscle gene expression** In the **Acute Study**, exercising before *versus* after nutrient provision increased the acute plasma glucose and serum insulin responses to food consumption (Figure 2A and 2B). The plasma glucose AUC was 6.70 [6.00 to 7.39] mmol·L^-1^·330 min^-1^ with exercise before nutrient provision *versus* 5.91 [5.33 to 6.50] mmol·L^-1^·330 min^-1^ with exercise after nutrient provision (*p<*0.01). The serum insulin AUC was 86.9 [48.5 to 125.2] pmol·L^-1^·330 min^-1^ with exercise before nutrient provision *versus* 55.3 [31.2 to 79.3] pmol·L^-1^·330 min^-1^ with exercise after nutrient provision (*p<*0.01). Exercise performed before *versus* after nutrient provision resulted in higher glycerol and non-esterified fatty acid (NEFA) concentrations during the exercise (Figure 2C and 2D).

**Figure 2.**
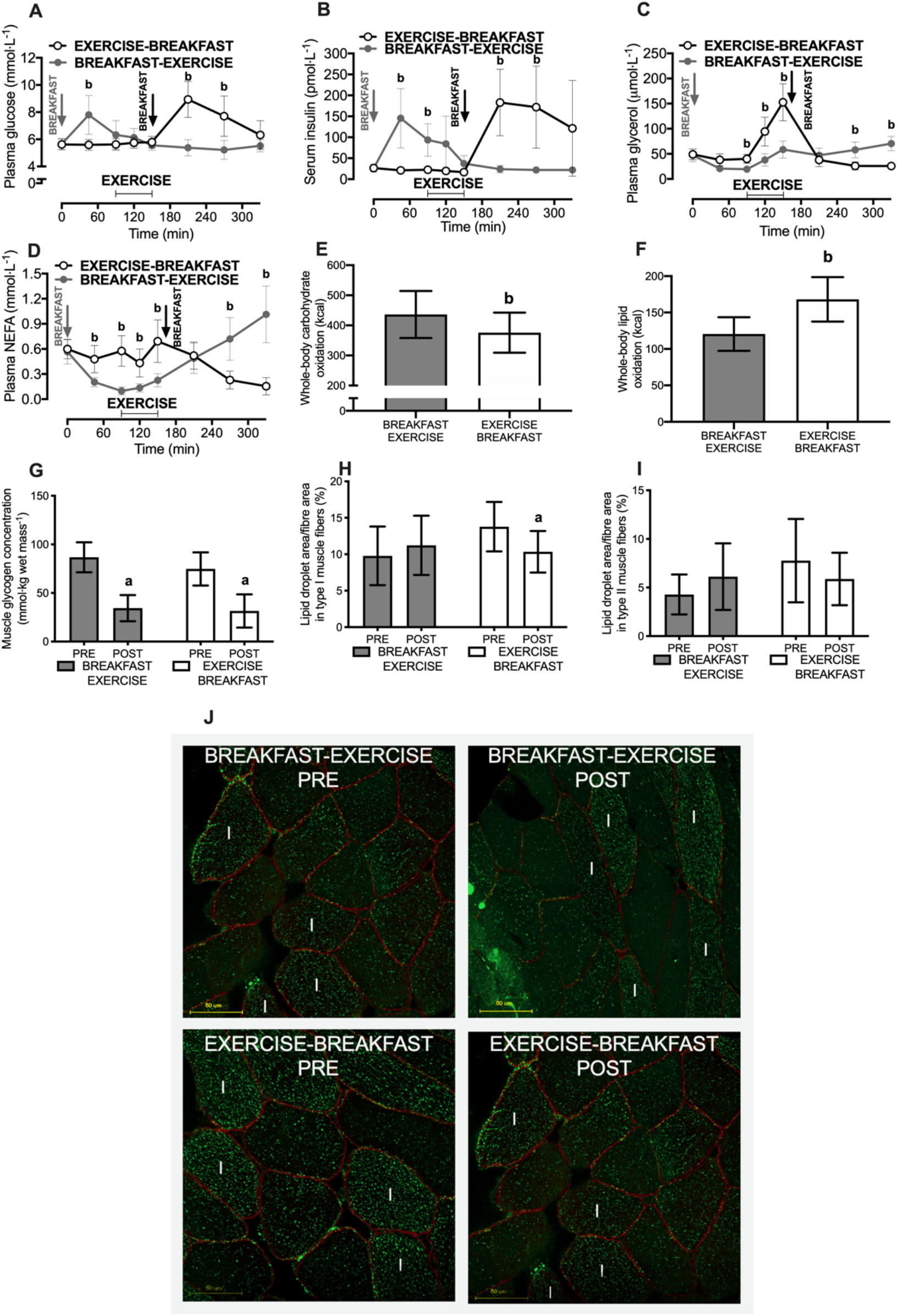
Plasma glucose (**A**), serum insulin (**B**), plasma glycerol (**C**) and plasma NEFA (**D**) concentrations and whole-body carbohydrate (**E**) and fat (**F**) utilization rates. Muscle was sampled pre-and immediately post-exercise (*vastus lateralis*) to assess mixed-muscle glycogen (**G**) and fiber-type specific intramuscular lipid (IMTG) utilization (**H & I**). Panel **J** is representative images from IMTG staining where IMTG (stained green) in combination with dystrophin (to identify the cell border and stained red) is shown from skeletal muscle samples of a representative participant for the breakfast-exercise and exercise-breakfast trials. White I shows type 1 fibers and all other fibers are assumed to be type II. Yellow bars are scale (50 μm). All data are presented as means ± 95% CI. For panels **A-F** *n*=12 men classified as overweight or obese, for panels **G**, **H and I** *n*=9. ^a^difference between PRE *versus* POST exercise; ^b^difference between BREAKFAST-EXERCISE *versus* EXERCISE-BREAKFAST (*p* < 0.05).

Nutrient provision before exercise potently altered whole-body metabolism, resulting in an increase in whole-body carbohydrate utilization (Figure 2E) and a decrease in whole-body lipid utilization (Figure 2F). In skeletal muscle, glycogen utilization during exercise (time effect, *p<*0.01) was independent of nutrient-exercise timing (time x trial interaction effect *p=*0.12; Figure 2G). However, the type I muscle fibre intramuscular triglyceride (IMTG) content was only reduced with exercise performed before nutrient provision (time x trial interaction: *p=*0.02; Figure 2H **and** 2J). A similar pattern was observed for the type II muscle fibre IMTG content (time x trial interaction: *p=*0.04; Figure 2I **and** 2J), although the reduction with exercise before nutrient provision did not achieve statistical significance after post-hoc corrections. Nonetheless, clear differences (both *p<*0.05) in the net changes in the IMTG content were observed in both fibre types with exercise before *versus* after nutrient provision (for type I: −3.44 [−1.61 to −5.26]% *versus* 1.44 [−1.46 to 4.34]% area covered by lipid staining and type II: −1.89 {-0.16 to −3.61]% *versus* 1.83 [0.50 to 3.17]% area covered by lipid staining for exercise before *versus* after nutrient provision, respectively).

Of the 34 selected genes that are implicated in metabolic adaptations to exercise, only 8 genes were altered by exercise, whereby *IRS-1* and *FATP1* were decreased post-exercise compared to baseline (*p<*0.05) and *IRS-2, PDK4, PGC1α, FATP4* and *ACSL1* were increased post-exercise compared to baseline (all *p<*0.05). However, only *PPARδ* was differentially expressed by nutrient-exercise timing and was higher with breakfast before *versus* after exercise (*p<*0.05; Figure 3).

**Figure 3.**
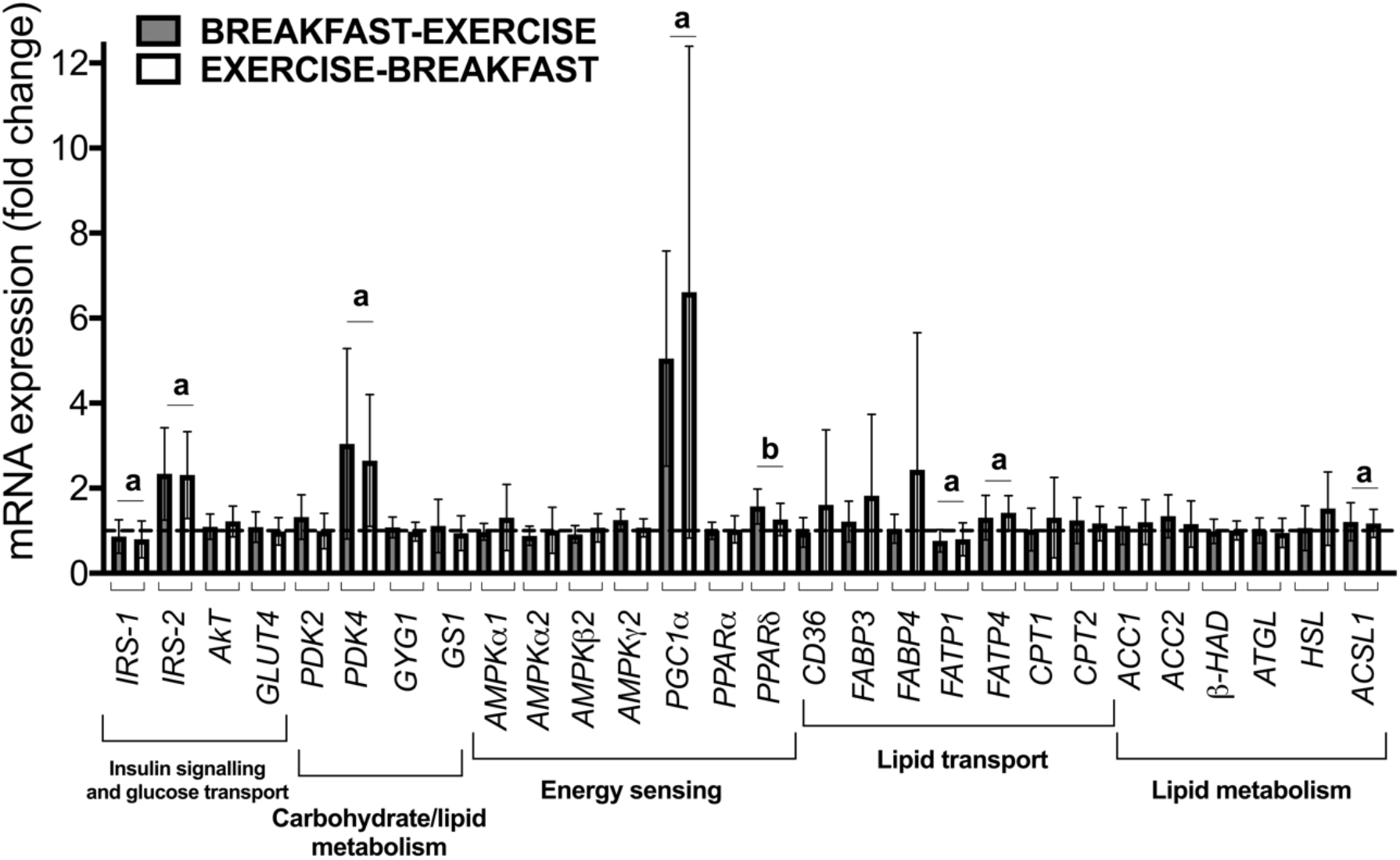
Skeletal muscle mRNA expression responses to a single bout of exercise before *versus* after nutrient provision (in the form of breakfast) in overweight men. *n*=8. Muscle was sample pre-and at 3 h post-exercise (*vastus lateralis*) to assess the intramuscular gene expression responses to exercise. ^a^effect of exercise; ^b^ difference between BREAKFAST-EXERCISE *versus* EXERCISE-BREAKFAST (*p* < 0.05).

### Exercise training before nutrient provision leads to sustained increases in lipid utilization

In the **Training Study** the compliance to the training was 100%, as all sessions were completed as prescribed. The average exercise intensity was 62 ± 5% V̇O_2_ peak in BR-EX and 62 ± 4% V̇O_2_ peak in EX-BR (*p=*0.98) and the heart rate (HR) response and average rating of perceived exertion (RPE) to the exercise training were 140 ± 13 *versus* 134 ± 8 beats·min^-1^ in BR-EX *versus* EX-BR (*p=*0.18) and 13 ± 1 au *versus* 13 ± 1 au (6-20 rating scale) in BR-EX *versus* EX-BR; (*p=*0.54), respectively.

In the **Training Study** rates of whole-body lipid utilization were around 2-fold higher with exercise before *versus* after nutrient provision and this difference between the conditions was sustained throughout the whole 6-week intervention (Figure 4A). As a consequence, regular exercise before (*versus* after) nutrient provision increased cumulative whole-body lipid utilization (during exercise) over a 6-week intervention, from 799 kcal [530 to 1069] in BR-EX to 1666 kcal [1260 to 2072] in EX-BR (*p<*0.01). This was accompanied by a decrease in rates of whole-body carbohydrate utilization during exercise (Figure 5A), as reflected by a decrease in the respiratory exchange ratio (group effect, *p<*0.01; Figure 5B). However, cumulative energy expenditure throughout the exercise intervention did not differ with exercise performed before *versus* after nutrient provision (Figure 5C; 7207 [6739 to 7676] kcal in BR-EX *versus* 6951 [6267 to 7635] kcal in EX-BR; *p=*0.48).

**Figure 4.**
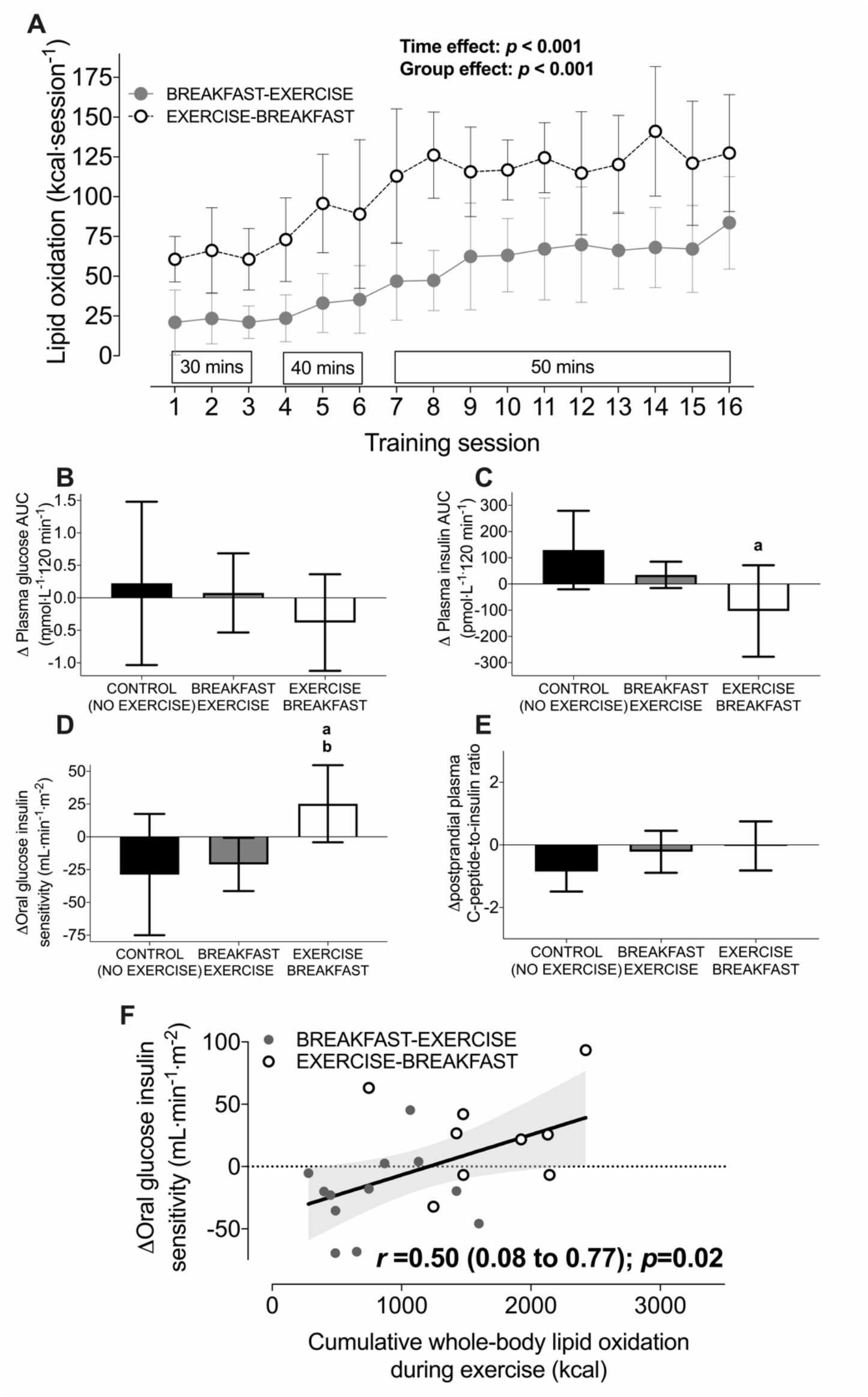
Whole-body lipid utilization during every exercise session in a 6-week training intervention (**A**), the change in the plasma glucose AUC (**B**), the change in the plasma insulin AUC (**C**), the change in the oral glucose insulin sensitivity index (OGIS; **D**) and the change in the postprandial plasma C-peptide: insulin ratio (**E**) in control (no-exercise), breakfast-exercise and exercise-breakfast groups. Panel **F** shows the Pearson correlation between changes in the OGIS index and cumulative lipid utilization throughout the exercise training intervention. All data are presented as means ± 95% CI. For control *n* = 9, for breakfast-exercise *n* = 12 and for exercise-breakfast *n* = 9 men classified as overweight or obese. The shaded grey area represents the 95% confidence bands for the regression line. ^a^difference between CONTROL *versus* EXERCISE-BREAKFAST; ^b^difference between BREAKFAST-EXERCISE *versus* EXERCISE-BREAKFAST (*p* < 0.05).

**Figure 5.**
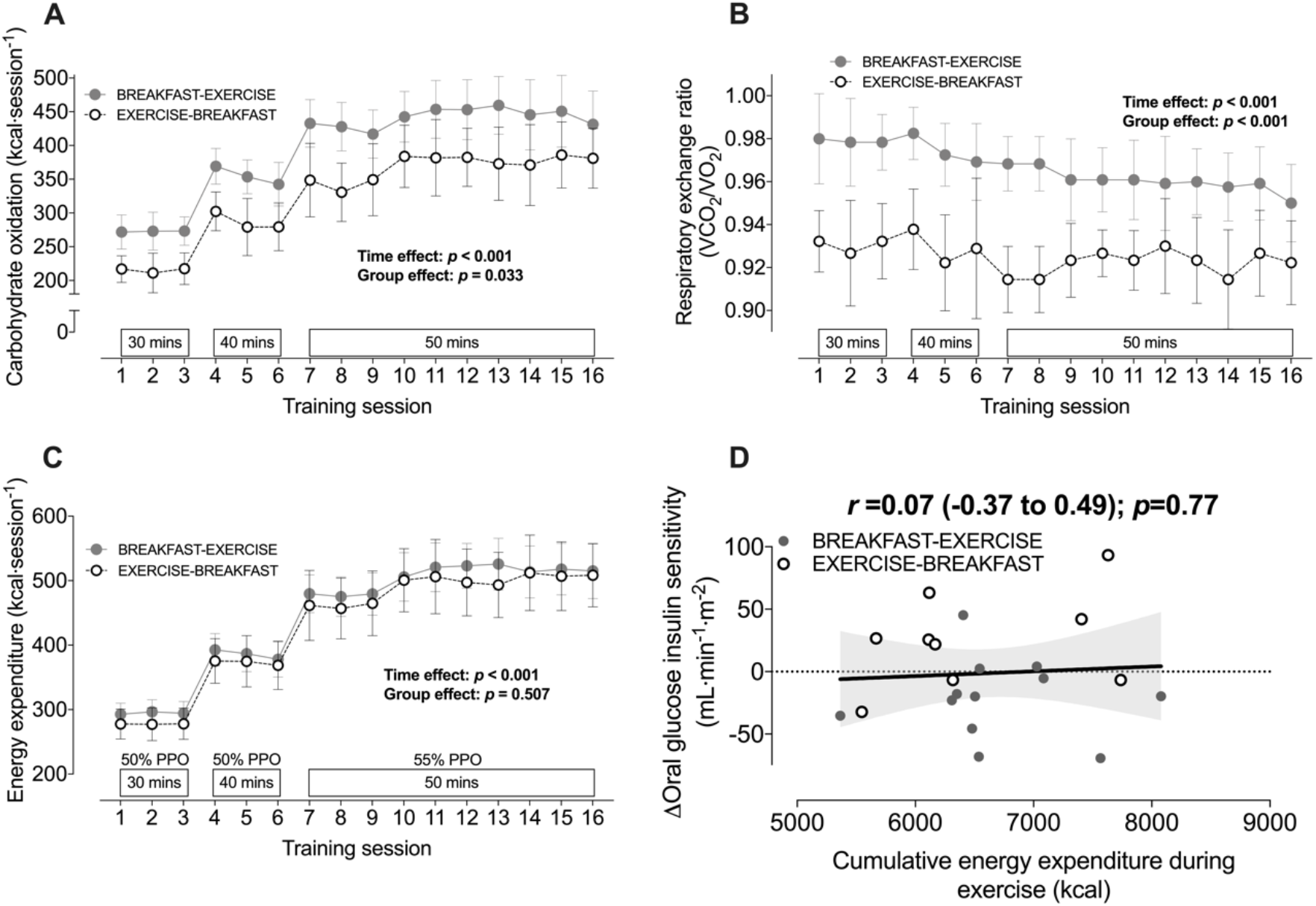
Energy expenditure (**A**), the respiratory exchange ratio (**B**) and whole-body carbohydrate utilisation rates (**C**) during every exercise session in a 6-week training intervention and a Pearson correlation between cumulative energy expenditure throughout the exercise training intervention with the changes in oral glucose insulin sensitivity (the OGIS index) with exercise before *versus* after nutrient intake (**D**). All data are presented as means ± 95% CI. For control *n* = 9, for breakfast-exercise *n* = 12 and for exercise-breakfast *n* = 9 men classified as overweight or obese. The shaded grey area represents the 95% confidence bands for the regression line.

### Exercise training before *versus* after nutrient provision increases an index of oral glucose insulin sensitivity

The oral glucose tolerance test-derived estimate of peripheral insulin sensitivity (the OGIS index; *p=*0.26), postprandial glycaemia (*p=*0.80) and postprandial insulinemia (*p=*0.30) were similar between groups pre-intervention (Table 3). The intervention-induced changes in postprandial glycaemia (time x group interaction, *p=*0.54; Figure 4B) and fasting blood lipid profiles were unaffected by nutrient-exercise timing (pre-and post-intervention data are shown in Tables 3 and 4). However, exercise training before, but not after nutrient intake reduced postprandial insulinemia (time x group interaction *p=*0.03; Figure 4C). Exercise training before *versus* after nutrient provision also increased the OGIS index (time x group interaction *p=*0.03; Figure 4D with pre-and post-intervention data in Table 3). The plasma C-peptide-to-insulin ratio was not differentially altered by nutrient-exercise timing (time x group interaction, *p=*0.12; Figure 4E). The change in the OGIS index in response to exercise training was positively and moderately correlated with cumulative lipid utilization during exercise throughout the intervention (Figure 4F) but not with cumulative energy expenditure (Figure 5C).

**Table 3.** Postprandial plasma metabolite concentrations for the control (CON; *n*=9), breakfast-exercise (BR-EX; *n*=12) and exercise-breakfast (EX-BR; *n*=9) groups.

|  | Pre-intervention | Post-intervention | *p*-value<br>(within-group<br>change) |
| --- | --- | --- | --- |
| CON glucose AUC ($\text{mmol}\cdot\text{L}^{-1}$) | 8.20 (1.36) | 8.43 (0.89) | 0.69 |
| BR-EX glucose AUC ($\text{mmol}\cdot\text{L}^{-1}$) | 8.26 (0.90) | 8.34 (1.35) | 0.78 |
| EX-BR glucose AUC ($\text{mmol}\cdot\text{L}^{-1}$) | 8.52 (1.05) | 8.14 (1.03) | 0.27 |
| CON insulin AUC ($\text{pmol}\cdot\text{L}^{-1}$) | 385 (221) | 514 (382) | 0.08 |
| BR-EX insulin AUC ($\text{pmol}\cdot\text{L}^{-1}$) | 268 (109) | 303 (135) | 0.16 |
| EX-BR insulin AUC ($\text{pmol}\cdot\text{L}^{-1}$) | 458 (441) | 355 (254) | 0.21 |
| CON C-peptide AUC ($\text{ng}\cdot\text{mL}^{-1}$) | 6.59 (2.57) | 7.32 (3.11) | 0.18 |
| BR-EX C-peptide AUC ($\text{ng}\cdot\text{mL}^{-1}$) | 5.25 (1.72) | 5.54 (1.87) | 0.34 |
| EX-BR C-peptide AUC ($\text{ng}\cdot\text{mL}^{-1}$) | 6.81 (4.40) | 6.03 (3.45) | 0.28 |
| CON NEFA AUC ($\text{mmol}\cdot\text{L}^{-1}$) | 0.17 (0.07) | 0.19 (0.07) | 0.52 |
| BR-EX NEFA AUC ($\text{mmol}\cdot\text{L}^{-1}$) | 0.20 (0.08) | 0.16 (0.06) | 0.08 |
| EX-BR NEFA AUC ($\text{mmol}\cdot\text{L}^{-1}$) | 0.14 (0.03) | 0.13 (0.04) | 0.48 |
| CON OGIS ($\text{mL}\cdot\text{min}^{-1}\cdot\text{m}^{-2}$) | 401 (39) | 372 (63) | 0.19 |
| BR-EX OGIS ($\text{mL}\cdot\text{min}^{-1}\cdot\text{m}^{-2}$) | 403 (31) | 380 (34) | 0.06 |
| EX-BR OGIS ($\text{mL}\cdot\text{min}^{-1}\cdot\text{m}^{-2}$) | 374 (56) | 399 (42) | 0.08 |

Data are means and (SD). Abbreviations: NEFA = non-esterified fatty acid, OGIS = oral glucose insulin sensitivity, AUC = time-averaged area under the curve for the OGTT (120 min).

**Table 4.** Fasting plasma metabolite concentrations for the control (CON; *n*=9), breakfast-exercise (BR-EX; *n*=12) and exercise-breakfast (EX-BR; *n*=9) groups.

| | Pre-intervention | Post-intervention | $\Delta$ from pre-intervention | time x group interaction |
| --- | --- | --- | --- | --- |
| CON glucose ($\text{mmol}\cdot\text{L}^{-1}$) | 5.39 (0.49) | 5.57 (0.66) | 0.17 (-0.32, 0.66) | F=1.413<br>$p=0.26$ |
| BR-EX glucose ($\text{mmol}\cdot\text{L}^{-1}$) | 5.48 (0.33) | 5.60 (0.47) | 0.12 (-0.30, 0.54) | |
| EX-BR glucose ($\text{mmol}\cdot\text{L}^{-1}$) | 5.72 (0.71) | 5.46 (0.72) | -0.27 (-0.67, 0.13) | |
| CON insulin ($\text{pmol}\cdot\text{L}^{-1}$) | 95 (121) | 81 (63) | -14 (-7, 8) | F=0.327<br>$p=0.72$ |
| BR-EX insulin ($\text{pmol}\cdot\text{L}^{-1}$) | 43 (23) | 43 (24) | 0 (-14, 15) | |
| EX-BR insulin ($\text{pmol}\cdot\text{L}^{-1}$) | 49 (43) | 47 (35) | -2 (-20, 14) | |
| CON HOMA-IR (au) | 0.35 (0.04) | 0.34 (0.03) | -0.01 (-0.02, 0.00) | F=0.458<br>$p=0.40$ |
| BR-EX HOMA-IR (au) | 0.37 (0.03) | 0.37 (0.04) | 0.00 (-0.01, 0.01) |  |
| EX-BR HOMA-IR (au) | 0.37 (0.04) | 0.37 (0.04) | 0.01 (-0.02, 0.04) |  |
| CON NEFA ($\text{mmol}\cdot\text{L}^{-1}$) | 0.36 (0.16) | 0.39 (0.11) | 0.03 (-0.10, 0.15) | F=1.021<br>$p=0.37$ |
| BR-EX NEFA ($\text{mmol}\cdot\text{L}^{-1}$) | 0.44 (0.15) | 0.39 (0.10) | -0.05 (-0.11, 0.15) | |
| EX-BR NEFA ($\text{mmol}\cdot\text{L}^{-1}$) | 0.34 (0.09) | 0.32 (0.10) | -0.02 (-0.09, 0.05) | |
| CON TAG ($\text{mmol}\cdot\text{L}^{-1}$) | 1.59 (0.77) | 2.05 (0.70) | 0.46 (-0.06, 0.98) | F= 5.967<br>$p<0.01$ |
| BR-EX TAG ($\text{mmol}\cdot\text{L}^{-1}$) | 1.34 (0.84) | 1.11 (0.42) | -0.24 (-0.55, 0.76) a | |
| EX-BR TAG ($\text{mmol}\cdot\text{L}^{-1}$) | 1.10 (0.31) | 0.88 (0.36) | -0.22 (-0.41, -0.03) b | |
| CON cholesterol ($\text{mmol}\cdot\text{L}^{-1}$) | 4.41 (1.23) | 4.59 (1.36) | 0.19 (-0.74, 1.12) | F=1.707<br>$p=0.20$ |
| BR-EX cholesterol ($\text{mmol}\cdot\text{L}^{-1}$) | 3.77 (1.34) | 3.72 (1.21) | -0.05 (-0.41, 0.30) | |
| EX-BR cholesterol ($\text{mmol}\cdot\text{L}^{-1}$) | 3.74 (0.82) | 3.24 (0.92) | -0.51 (-0.95, -0.07) | |
| CON HDL cholesterol ($\text{mmol}\cdot\text{L}^{-1}$) | 0.84 (0.17) | 0.86 (0.20) | 0.02 (-0.14, 0.18) | F=1.634<br>$p=0.21$ |
| BR-EX HDL cholesterol ($\text{mmol}\cdot\text{L}^{-1}$) | 0.82 (0.31) | 0.84 (0.33) | 0.02 (-0.05, 0.10) | |
| EX-BR HDL cholesterol ($\text{mmol}\cdot\text{L}^{-1}$) | 0.90 (0.23) | 0.81 (0.26) | -0.09 (-0.19, 0.01) | |
| CON LDL cholesterol ($\text{mmol}\cdot\text{L}^{-1}$) | 3.36 (1.33) | 3.56 (1.36) | 0.20 (-0.50, 0.89) | F=2.110<br>$p=0.14$ |
| BR-EX LDL cholesterol ($\text{mmol}\cdot\text{L}^{-1}$) | 2.71 (0.97) | 2.70 (0.99) | -0.01 (-0.31, 0.28) | |
| EX-BR LDL cholesterol ($\text{mmol}\cdot\text{L}^{-1}$) | 2.72 (0.57) | 2.31 (0.58) | -0.41 (-0.80, -0.02) | |

Data are means and (SD) except for change scores which are means and (95% CI). Abbreviations: HOMA-IR = the homeostatic model of insulin resistance, NEFA = non-esterified fatty acid, TAG = triglyceride, HDL = high density lipoprotein, LDL = low density lipoprotein. (a) = difference in change from pre- to post-intervention for CON *versus* BR-EX and (b) CON *versus* EX-BR with $p<0.05$ .

### Nutrient-exercise timing does not differentially alter body composition or oxidative capacity

Exercise before *versus* after nutrient provision resulted in comparable changes in body mass (time x group interaction, *p=*0.97; Figure 6A), a marker of central adiposity (the waist to hip ratio; time x group interaction, *p=*0.17, Figure 6B), and the peak capacity for whole-body lipid utilization (time x group interaction, *p=*0.14; Figure 6C). Exercise training increased V̇O_2_ peak by ∼ 3 mL×kg×min^-1^ relative to a no-exercise control (CON) group (time x group interaction, *p=*0.01) but the magnitude of this increase in cardiorespiratory fitness was unaffected by nutrient exercise timing (*p=*0.54 with breakfast-exercise *versus* exercise-breakfast). Self-reported daily energy intake was unaffected by exercise or nutrient-exercise timing (time x group interaction, *p=*0.38; Table 5), and although daily energy expenditure was increased in the exercise groups *versus* control group (time x group interaction, *p=*0.01; Table 5), this increase was unaffected by nutrient-exercise timing (*p=*0.38).

**Figure 6.**
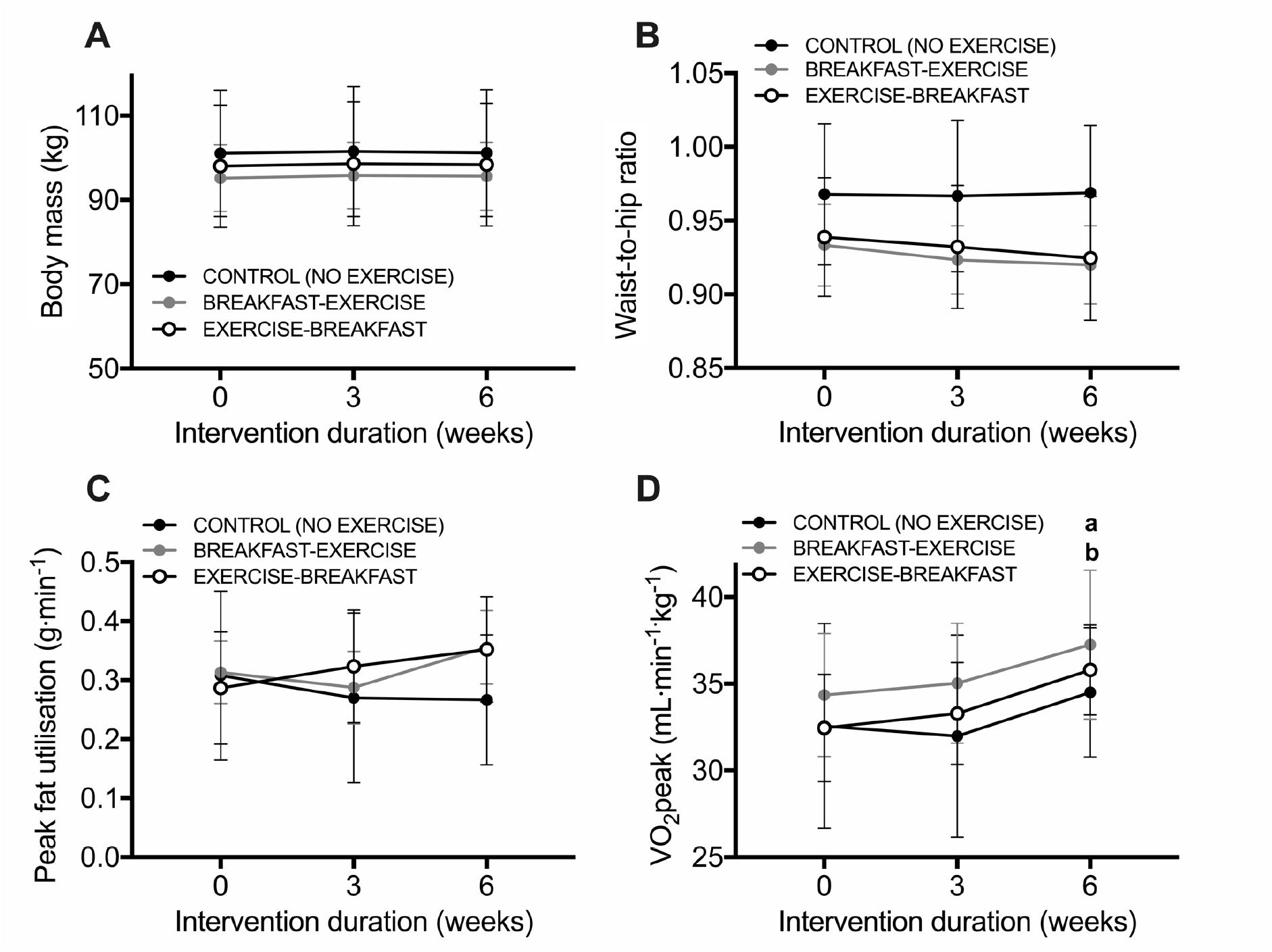
Body mass (**A**), the waist-to-hip ratio (**B**), whole-body oxidative capacity (VO_2_peak; **C**) and peak fat utilization rates during an incremental exercise test (**D**) at baseline, week 3 and week 6 of an intervention in control (no-exercise), breakfast-exercise and exercise-breakfast groups. All data are presented as means ± 95% CI. For control *n* = 9, for breakfast-exercise *n* = 12 and for exercise-breakfast *n* = 9 men classified as overweight or obese ^a^difference between CONTROL *versus* BREAKFAST-EXERCISE; ^b^difference between CONTROL *versus* EXERCISE-BREAKFAST (*p* < 0.05).

**Table 5.** Components of daily energy intake and daily energy expenditure pre-and post-intervention for the control (CON; n=9 [except for PAEE where *n*=8]), breakfast before exercise (BR-EX; n=12 [except for PAEE where *n*=10]) and exercise before breakfast (EX-BR; n=9) groups.

| | Pre-intervention | Post-intervention | $\Delta$ from pre-intervention | time x group interaction |
| --- | --- | --- | --- | --- |
| CON CHO intake (kcal·d<sup>-1</sup>) | 1180 (357) | 1424 (600) | 244 (-29, 517) | F=0.977<br>p=0.39 |
| BR-EX CHO intake (kcal·d<sup>-1</sup>) | 1171 (317) | 1272 (220) | 101 (-51, 252) |  |
| EX-BR CHO intake (kcal·d<sup>-1</sup>) | 1133 (307) | 1214 (234) | 81 (-91, 252) |  |
| CON FAT intake (kcal·d<sup>-1</sup>) | 1148 (326) | 1045 (311) | -104 (-257, 49) | F=0.347<br>p=0.71 |
| BR-EX FAT intake (kcal·d<sup>-1</sup>) | 1132 (307) | 1091 (238) | -41 (-156, 75) |  |
| EX-BR FAT intake (kcal·d<sup>-1</sup>) | 987 (320) | 882 (172) | -105 (-289, 79) |  |
| CON PRO intake (kcal·d<sup>-1</sup>) | 429 (97) | 376 (89) | -52 (-89, 15) | F=1.517<br>p=0.24 |
| BR-EX PRO intake (kcal·d<sup>-1</sup>) | 511 (122) | 446 (59) | -65 (-118, -11) |  |
| EX-BR PRO intake (kcal·d<sup>-1</sup>) | 406 (97) | 399 (117) | -7 (-75, 62) |  |
| CON ALC intake (kcal·d<sup>-1</sup>) | 72 (95) | 75 (154) | 3 (-145, 152) | F=1.115<br>p=0.34 |
| BR-EX ALC intake (kcal·d<sup>-1</sup>) | 111 (118) | 69 (72) | -42, (-101, 17) |  |
| EX-BR ALC intake (kcal·d<sup>-1</sup>) | 185 (168) | 90 (131) | -94 (-191, 2) |  |
| CON RMR (kcal·d<sup>-1</sup>) | 1997 (225) | 2021 (232) | 24 (-56, 104) | F=1.791<br>p=0.19 |
| BR-EX RMR (kcal·d<sup>-1</sup>) | 1964 (167) | 2074 (208) | 110 (45, 174) |  |
| EX-BR RMR (kcal·d<sup>-1</sup>) | 1899 (228) | 1967 (194) | 68 (-13, 148) |  |
| CON TEF (kcal·d<sup>-1</sup>) | 283 (61) | 292 (85) | 9 (-29, 48) | F=0.982<br>p= 0.38 |
| BR-EX TEF (kcal·d<sup>-1</sup>) | 292 (50) | 288 (42) | -4 (-19, 10) |  |
| EX-BR TEF (kcal·d<sup>-1</sup>) | 271 (55) | 258 (44) | -13 (-29, 4) |  |
| CON PAEE (kcal·d<sup>-1</sup>) | 1144 (298) | 1077 (327) | -67 (-215, 81) | F=7.044<br>p<0.01 |
| BR-EX PAEE (kcal·d<sup>-1</sup>) | 986 (264) | 1190 (343) | 204 (56, 352) a |  |
| EX-BR PAEE (kcal·d<sup>-1</sup>) | 1006 (147) | 1357 (324) | 351 (126, 576) b |  |

Data are means and (SD) except for change scores which are means and (95% CI). Abbreviations: CHO = carbohydrate, PRO = protein, ALC = alcohol, RMR = resting metabolic rate, TEF = thermic effect of feeding, PAEE = physical activity energy expenditure. (a) = difference in change from pre- to post-intervention for CON *versus* BR-EX and (b) CON *versus* EX-BR with p<0.05.

**Table 6.**
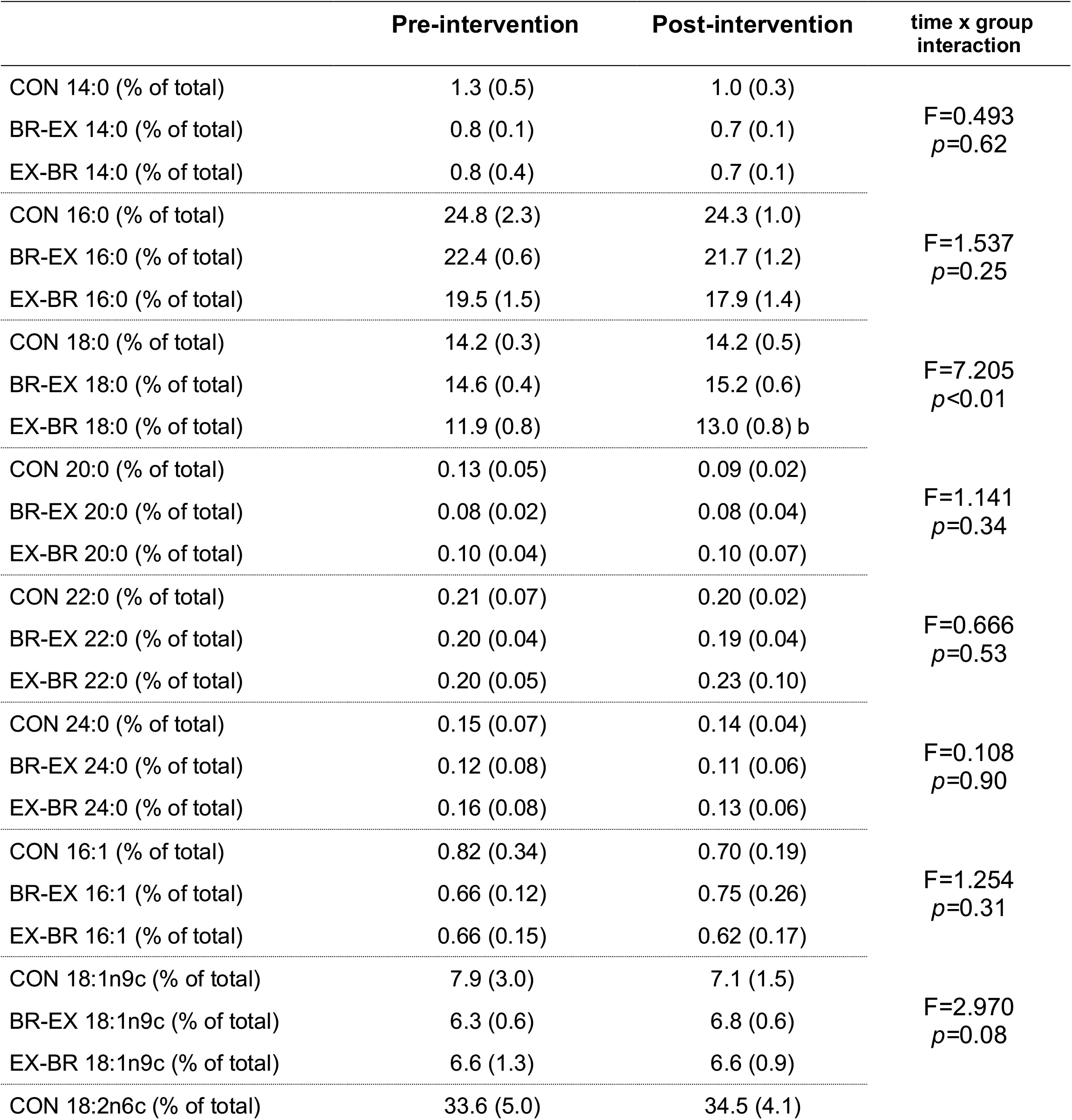
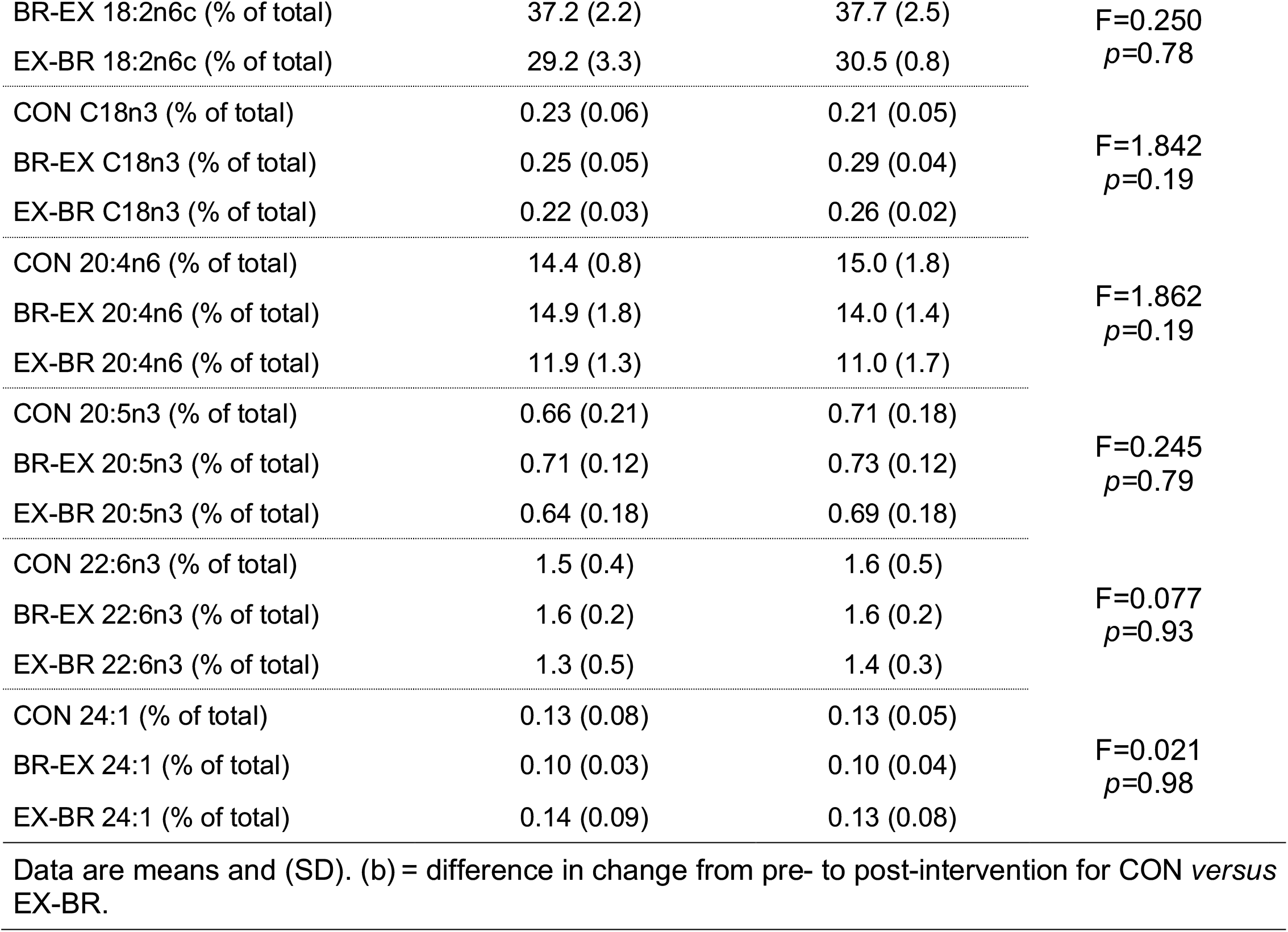
Skeletal muscle phospholipid composition in the control (CON; *n*=6), breakfast-exercise (BR-EX; *n*=8) and exercise-breakfast (EX-BR; *n*=5) groups.

### Exercise training before nutrient provision increases phospholipid remodeling

There was a significant effect on global skeletal muscle remodeling with exercise before *versus* after nutrient ingestion as quantified by the sum of changes in the fatty acid content of all phospholipid species (*p=*0.01; Figure 7A). Nutrient provision prior to exercise prevented this exercise-induced increase in skeletal muscle phospholipid remodeling as the sum of changes was not different to a non-exercise control group (*p=*0.41). No clear time x group interaction effects were determined for any of the measured fatty acid species, except for the proportion of 18:0, which increased with exercise before nutrient provision compared to the control group (Table 6). The change in the overall saturated fatty acid content of skeletal muscle phospholipids was moderately and positively correlated with changes in postprandial insulinemia and the relationship was robust to the exclusion of any single data point (Figure 7B).

**Figure 7.**
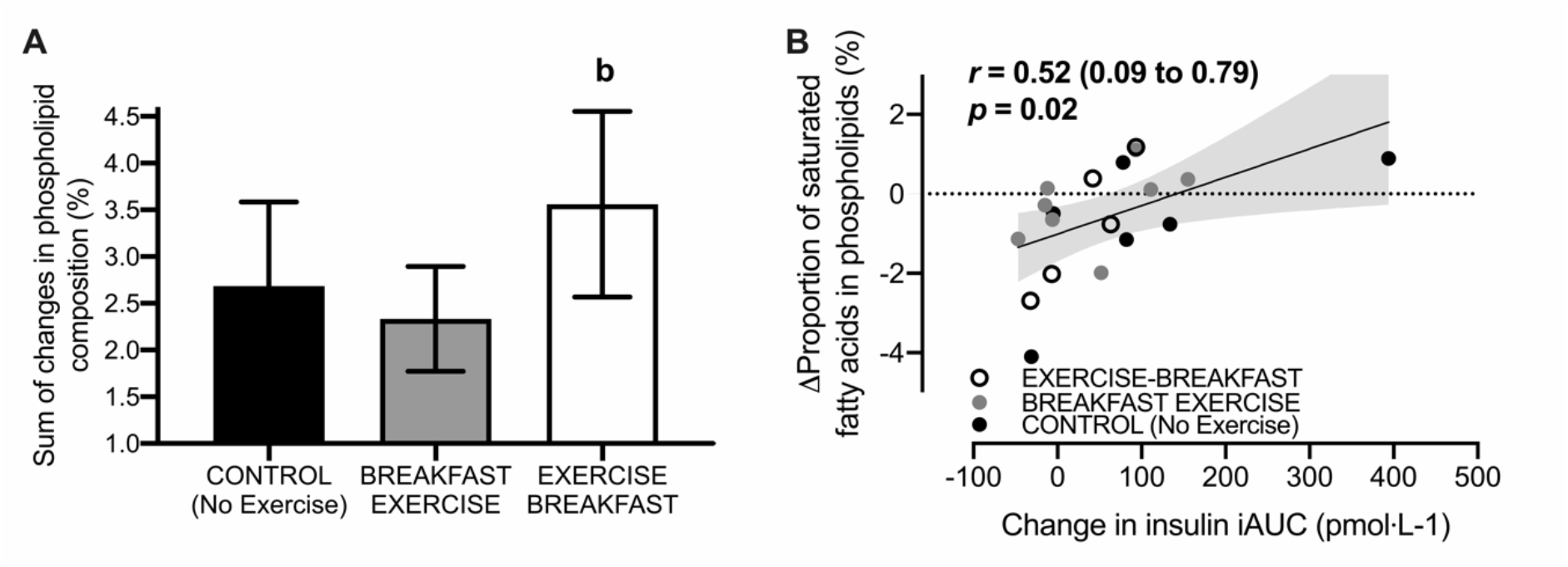
Pre-to-post intervention changes in the sum of all changes in the fatty acid content of phospholipid species (**A**) and a Pearson correlation between postprandial insulinemia with the change in the proportion of saturated fatty acids in skeletal muscle phospholipids (**B**). All data are presented as means ± 95% CI. For control *n* = 6, for breakfast-exercise *n* = 9 and for exercise-breakfast *n* = 5 men classified as overweight or obese. The shaded area represents the 95% confidence bands for the regression line. ^b^difference between BREAKFAST-EXERCISE *versus* EXERCISE-BREAKFAST (*p* < 0.05).

### Exercise training before nutrient provision augments intramuscular adaptations

Skeletal muscle AMPK protein levels increased ∼3-fold with exercise training performed before-but not after-nutrient provision *versus* a no-exercise control group (Figure 8A). However, these increases did not translate into differential changes in proteins including CD36 and CPT-1 which are involved in fatty acid transport in skeletal muscle [both *p>*0.05; data available online (51)], or markers of mitochondrial oxidative capacity, including the protein levels of the OXPHOS complexes [all *p>*0.05; data available online (51)] or citrate synthase activity (change from baseline: −2.1 μmol×min^-1^×mg of protein^-1^ [-12.9 to 8.7] in CON, 7.6 μmol×min^-1^×mg of protein^-1^ [-1.2 to 16.4] in BR-EX and 6.5 μmol×min^-1^×mg of protein^-1^ [0.2 to 12.8] in EX-BR; *p>*0.05). There were also no differential changes in the content of insulin signaling proteins such as Akt2 or AS160 in response to nutrient-exercise timing (*p>*0.05; Figure 8B). However, there was a ∼2-fold increase in skeletal muscle GLUT4 protein levels with exercise training performed before (*p=*0.04), but not after nutrient provision (*p=*0.58) *versus* a non-exercise control group (Figure 8A). There was also an increase in the protein levels of the CHC22 clathrin isoform and its associated adaptor protein (GGA2) relative to the CHC17 clathrin isoform, with exercise before *versus* after nutrient provision (both *p<*0.05; Figure 8B). When we examined the CHC22 isoform alone [data not shown but available online (51)]we noted baseline differences which may have confounded the interpretation of these fold-changes due to regression to the mean. We thus present the CHC22/CHC17 ratio (Figure 8B) to reflect GLUT4-associated clathrin-mediated membrane traffic relative to total clathrin-mediated membrane traffic.

**Figure 8.**
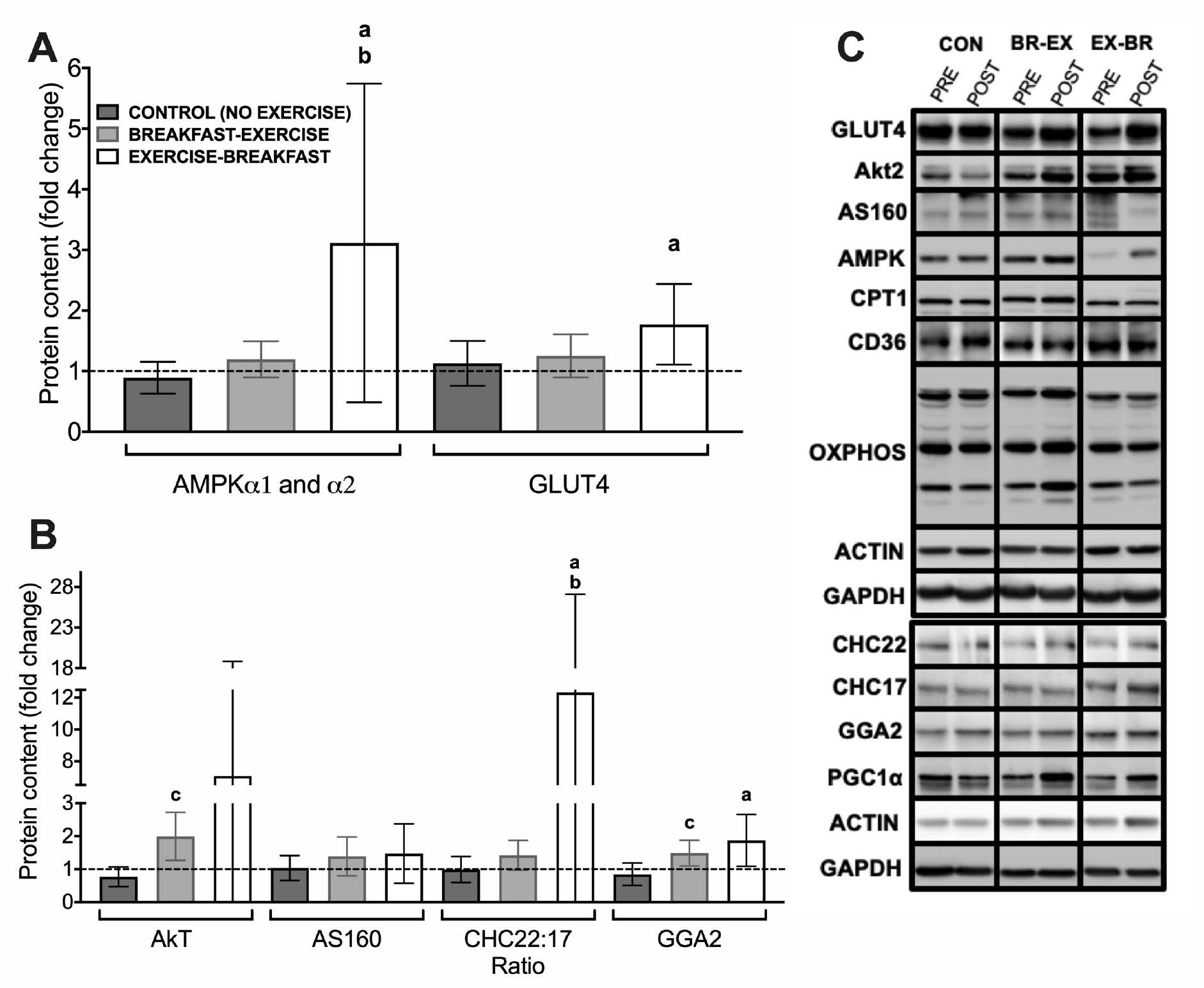
Pre-to-post intervention changes in the levels of energy-sensing proteins and proteins involved in insulin-sensitive GLUT4 trafficking in skeletal muscle (**A** and **B**). Representative immunoblots are shown (**C**) for each protein (including those reported in text but not shown in this figure) from the same representative participant as well as the loading controls used. All data are presented as means ± 95% CI and the dotted horizontal line represents the baseline (pre-intervention) values. For control *n* = 6, for breakfast-exercise *n* = 9 and for exercise-breakfast *n* = 5 men classified as overweight or obese. ^a^difference between CONTROL *versus* EXERCISE-BREAKFAST; ^b^difference between BREAKFAST-EXERCISE *versus* EXERCISE-BREAKFAST; ^c^difference between CONTROL *versus* BREAKFAST-EXERCISE (*p* < 0.05).

## DISCUSSION

This is the first study to investigate the effect of nutrient-exercise interactions on key aspects of metabolic health in people classified as overweight or obese. We found that a single exercise bout performed before, but not after, nutrient provision increased whole body and skeletal muscle lipid utilization. We then used a 6-week training program to reveal sustained, 2-fold increases in lipid utilization that were maintained throughout 6 weeks of exercise training performed before *versus* after nutrient provision. An oral glucose tolerance test-derived estimate of peripheral insulin sensitivity (the OGIS index) increased with exercise training before *versus* after nutrient provision and this was associated with increased lipid utilization during the exercise training intervention. Exercise training prior to nutrient provision also augmented remodeling of phospholipids and increased the levels of energy sensing (i.e. AMPK) and glucose transport proteins (i.e. GLUT4) in exercised skeletal muscle. These results indicate that nutrient-exercise timing modulates training responsiveness in overweight men and link lipid utilization during exercise to exercise-training-induced changes in aspects of metabolic health.

First, we showed that a single bout of exercise performed before *versus* after nutrient intake increased whole-body lipid utilization. A blunting of intramuscular triglyceride (IMTG) utilization has been shown in type I fibers of lean, healthy men in response to carbohydrate ingestion before and during exercise, compared to exercise in the fasted-state (22). Here, we demonstrated for the first time that exercise before *versus* after breakfast consumption increases net IMTG utilization in men classified as overweight or obese. Whilst the authors do acknowledge that absolute IMTG content may have been underestimated due to the analytical procedures used to estimate IMTG (i.e. use of Triton-X100 detergent, overnight drying of mounting medium), all samples were treated consistently. We also showed that net skeletal muscle glycogen utilization and acute skeletal muscle mRNA responses were largely unaffected by the same exercise performed before *versus* after breakfast. This is important, because muscle glycogen availability can alter muscle adaptations to training (27). Lower muscle glycogen concentrations are therefore unlikely to have driven the training responses we observed in the training study with the present method of nutrient-exercise timing.

Altering substrate availability can also drive adaptive responses to exercise partly by modulating acute mRNA expression in exercised skeletal muscle (52). However, in the present study, only one measured gene was differentially expressed in response to exercise before *versus* after nutrient provision. Specifically, we observed less of an exercise-induced increase in skeletal muscle *PPARδ* expression with exercise before *versus* after nutrient provision, which is surprising given that *PPARδ* has been implicated in adaptations relating to oxidative capacity and lipid utilization (53). However, previous research has also shown no differential increase in *PPARδ* expression in skeletal muscle when exercise was performed with carbohydrate consumption before and during exercise *versus* in the fasted state (54). The different response observed in the present study might be because we assessed the effect of nutrient-exercise timing (*i.e.* nutrient provision before *versus* after exercise) rather than the omission *versus* ingestion of nutrients. This suggests that inferences cannot necessarily be extrapolated from studies assessing the effects of nutrient ingestion *versus* nutrient omission, to inform responses to models of nutrient-exercise *timing*.

In the training study, we then showed that the acute increases in whole-body lipid utilization during a single bout of exercise performed before *versus* after nutrient intake were sustained throughout 6-weeks of exercise training. Moreover, only exercise training performed before nutrient intake reduced postprandial insulinemia and increased the oral glucose tolerance test-derived estimate of peripheral insulin sensitivity (i.e. the OGIS index). As the plasma C-peptide-to-insulin ratio was not differentially altered by nutrient-exercise timing, the reduction in postprandial insulinemia with exercise performed before *versus* after nutrient ingestion is likely to be due to a reduction in insulin secretion rather than an increase in hepatic insulin extraction (55). It should also be noted that difference between the exercise groups for the change in the OGIS index was also broadly equivalent to the difference between individuals classified as having a healthy phenotype compared to individuals with impaired glucose tolerance (56).

Exercise training before *versus* after nutrient provision also resulted in augmented phospholipid remodeling in skeletal muscle. Moreover, the change in the saturated fatty acid content of skeletal muscle phospholipids with exercise correlated with the change in postprandial insulinemia. This supports prior observations that a higher proportion of saturated fatty acids in skeletal muscle phospholipids negatively correlates with insulin sensitivity (57). Single-leg exercise training has been used to show increased polyunsaturated fatty acid content of skeletal muscle phospholipids in an exercised *versus* non-exercised leg (58). Since that change was independent of dietary intake, the reduction in the saturated fatty content of phospholipids was likely due to a preferential upregulation of saturated fatty acid oxidation as a result of the higher energy expenditure (59, 60). However, because this previous work involved changes in energy expenditure across experimental conditions, the role of lipid utilization independent of energy expenditure on phospholipid remodeling could not be explored. Here, we showed that skeletal muscle remodeling was increased with exercise performed before *versus* after nutrient provision, presumably due to increased lipid utilization in that condition.

AMPK is also nutrient sensitive and contributes to regulation of fatty acid utilization (61), mitochondrial biogenesis (62) and the expression of proteins involved in skeletal muscle glucose uptake, including GLUT4 and AS160 (63–65), which are key players in whole-body insulin sensitivity (66). We observed greater increases in the protein content of AMPK in skeletal muscle with exercise training before *versus* after nutrient intake. The increase in the GLUT4 content of skeletal muscle we observed with exercise before nutrient provision may be explained by this heightened AMPK response and, in turn, may have contributed to increases in the OGIS index following exercise training before *versus* after nutrient provision (67). Skeletal muscle AMPK can be activated by increased fatty acid availability, independent of muscle glycogen and AMP concentrations (68). Muscle glycogen utilization can modulate *AMPK* and *GLUT4* mRNA expression with different exercise models (66). However, since we observed no difference in muscle glycogen utilization with altered nutrient-exercising timing in the acute study, the change in the GLUT4 content with exercise training before *versus* after breakfast is likely to be attributable to repeated increases in fatty acid availability, potentially through increases in the skeletal muscle AMPK content. The AMPK antibody we used detects both isoforms of the catalytic subunits of AMPK (AMPKα1 and α2). In human skeletal muscle, three different complexes have been described [α2β2γ1, α2β2γ3, and α1β2γ1; (69)] and our antibody therefore captured all complexes. Accordingly, we cannot speculate whether a specific heterotrimeric AMPK complex is predominately contributing to the increase in AMPK content that we report. As such, the effect of nutrient-exercise timing on AMPK activation warrants continued investigation.

The correct targeting and sequestration of GLUT4 into its intracellular insulin-responsive compartments is also important for insulin sensitivity in skeletal muscle (70, 71). Clathrin heavy chain isoform 22 (CHC22) plays a specialized role in regulating GLUT4 sequestration in human skeletal muscle (72), protecting GLUT4 from degradation (73) and making it more available for insulin-stimulated release. We showed an increase in CHC22 protein levels in exercised muscle (relative to the exercise effects on CHC17 protein levels) with exercise before *versus* after nutrient provision. As the cognate clathrin CHC17 plays a widespread membrane traffic role in many tissues, CHC17 levels provide a benchmark for general membrane traffic changes compared to those in the GLUT4 pathway (74). The relative increase in CHC22 levels we observed thus suggests that exercise before nutrient provision not only augments GLUT4 protein levels, but potentially also the machinery necessary for the appropriate sequestration and targeting of GLUT4 to its insulin-responsive compartment. This may lead to improved GLUT4 translocation and contribute to the increases in the OGIS index we observed with exercise training before *versus* after nutrient intake. However, the CHC22 results reported here should be interpreted cautiously due to the relatively small sample size and variability in the individual CHC22 responses. Further work is therefore needed to investigate nutrient-exercise interactions and their effect on CHC22 levels. In addition, the remodeling of skeletal muscle phospholipids could have contributed to the ability of GLUT4 to fuse to the muscle-plasma membrane via less rigid arrays of phospholipid molecules in plasma membranes (75).

The greater increase in AMPK content we observed with exercise before nutrient provision did not further augment measured markers of mitochondrial biogenesis in skeletal muscle in response to exercise training in overweight men. This is in contrast to prior work demonstrating that carbohydrate ingestion before and during exercise suppresses exercise-induced increases in the content of proteins in skeletal muscle involved in fatty acid transport and oxidation (26). This further highlights that the model of nutrient-exercise timing that we employed (breakfast consumption before *versus* after exercise) might be distinct from other types of nutrient timing. Although changes in skeletal muscle mitochondrial content and/or oxidative capacity may be involved in regulating insulin sensitivity (76), the lack of differential response with exercise before *versus* after nutrition provision in this study suggests that these factors are unlikely to explain the changes in the OGIS index with the current model of nutrient-exercise timing employed (i.e. exercise before versus after breakfast). It is also interesting that the intramuscular adaptations and changes in the OGIS index that we observed occurred in the presence of similar changes in body composition, self-reported daily dietary intake and total daily energy expenditure with altered nutrient-exercise timing. Notwithstanding other factors that may have contributed to the increases in oral glucose insulin sensitivity with exercise before *versus* after nutrient ingestion, this highlights lipid metabolism as a potentially important mechanism explaining the improvement in OGIS with regular exercise performed before *versus* after breakfast.

It should also be noted that the responses observed for OGIS were an interaction between groups, and thus the response to exercise before nutrient provision is an increase relative to the non-exercise control group and the exercise after nutrient intake group. Accordingly, these data may be specific to high-carbohydrate provision and although this is typical of breakfasts in developed countries, it remains to be seen whether lower-carbohydrate meals produce similar effects. Potential limitations in our work also include the absence of a non-exercise fasting group, which would have allowed us to explore the role of extended morning fasting *per se* in the training study. However, our prior work has already shown that extended morning fasting in an absence of exercise may impair insulin sensitivity and increase postprandial insulinemia in obese humans (77).

To summarize, the present data are the first to show that exercise training before *versus* after carbohydrate (i.e. breakfast) consumption affects responsiveness to exercise training in men classified as overweight or obese, including greater remodeling of skeletal muscle phospholipids, adaptations of proteins involved in nutrient sensing and glucose transport in skeletal muscle, and increases in and index of oral glucose insulin sensitivity. These data suggest that exercising in a fasted state can augment the adaptive response to exercise, without the need to increase the volume, intensity, or perception of effort of exercise. These responses may be linked to the acute increases in lipid utilization during every bout of exercise performed in the fasted-*versus* the fed-state (a difference that is sustained throughout a period of training over 6-weeks). These findings therefore have implications for future research and clinical practice. For example, exercise training studies should account for nutrient-exercise timing if aspects of metabolic control are an outcome measure. Secondly, to increase lipid utilization and oral glucose insulin sensitivity with training, endurance-type exercise should be performed before *versus* after nutrient intake (i.e. in the fasted state).

## ADDITIONAL INFORMATION

### Data Availability

Raw data are available as online supporting information (https://researchportal.bath.ac.uk/en/datasets/).

## Acknowledgments

The authors thank Russell Davies, Esther Punter, Emily Fallon, Josh Dominy and Lauren Davey for assisting and supervising some of the exercise training sessions, Marine Camus for technical advice on Western blotting and Laura Wood for assisting with Western blotting. We also thank all those who participated in the studies for their time and commitment.

